# Mechanism of TAX1BP1 recruitment in aggrephagy to switch from cargo collection to sequestration

**DOI:** 10.1101/2024.05.17.594671

**Authors:** Bernd Bauer, Jonas Idinger, Martina Schuschnig, Luca Ferrari, Sascha Martens

**Affiliations:** Max Perutz Labs, Vienna Biocenter Campus (VBC), Dr.-Bohr-Gasse 9, 1030, Vienna, Austria; University of Vienna, Max Perutz Labs, Department of Biochemistry and Cell Biology, Dr.-Bohr-Gasse 9, 1030, Vienna, Austria; Vienna Biocenter PhD Program, a Doctoral School of the University of Vienna and the Medical University of Vienna, A-1030 Vienna, Austria

**Author notes:** To whom correspondance should be adressed.

**Keywords:** Quality control, selective autophagy, aggrephagy p62, NBR1, TAX1BP1

## Abstract

Autophagy mediates the degradation of harmful material within lysosomes. In aggrephagy, the pathway mediating the degradation of aggregated, ubiquitinated proteins, this cargo material is collected in larger condensates prior to its sequestration by autophagosomes. In this process, SQSTM1/p62 and NBR1 drive cargo condensation, while TAX1BP1, which binds to NBR1 recruits the autophagy machinery to facilitate autophagosome biogenesis at the condensates. The mechanistic basis for the TAX1BP1 mediated switch from cargo collection to its sequestration is unclear. Here we show that TAX1BP1 is not a constitutive component of the condensates. Its recruitment correlates with the induction of autophagosome biogenesis. TAX1BP1 is sufficient to recruit the TBK1 kinase via the SINTBAD adapter. We define the NBR1 - TAX1BP1 binding site, which is adjacent to the GABARAP/LC3 interaction site and demonstrate that the recruitment of TAX1BP1 to cargo mimetics can be enhanced by an increased ubiquitin load. Our study suggests that autophagosome biogenesis is initiated once sufficient cargo is collected in the condensates.

## Introduction

Maintaining a healthy proteome is essential for cellular homeostasis and the survival of the organism. Cells have evolved numerous overlapping pathways to ensure that misfolded proteins are either refolded or swiftly ubiquitinated and removed by degradation (Chen *et al*, 2011). Apart from the ubiquitin-proteasome system, macroautophagy (hereafter autophagy) is a major pathway for the degradation of misfolded, ubiquitinated proteins (Jayaraj *et al*, 2020). The selective removal of these proteins is termed aggrephagy (Lamark & Johansen, 2021). In aggrephagy, ubiquitinated cargo proteins are collected in larger condensates and subsequently sequestered within autophagosomes, which form *de novo* around the condensates by the concerted action of the autophagy machinery (Bauer *et al*, 2023; Melia *et al*, 2020; Nishimura & Tooze, 2020). Defects in this process are associated with various neurodegenerative diseases (Yamamoto *et al*, 2023).

Cargo receptors orchestrate selective autophagy by binding the cargo material, recruiting the autophagy machinery for autophagosome biogenesis and subsequently linking the cargo to the GABARAP and LC3 decorated autophagosomal membrane (Adriaenssens *et al*, 2022). At least three cargo receptors cooperate in the degradation of ubiquitinated proteins by aggrephagy. p62 forms polymers and drives the formation of condensates with ubiquitinated cargoes through its UBA mediated interaction with the ubiquitin chains (Bjorkoy *et al*, 2005; Ciuffa *et al*, 2015; Kageyama *et al*, 2021; Sun *et al*, 2018; Turco *et al*, 2021; Zaffagnini *et al*, 2018). NBR1 binds to p62 via its N-terminal PB1 domain and assists p62 in cargo condensation by contributing a high affinity UBA domain to the complex (Kirkin *et al*, 2009; Lamark *et al*, 2003; Turco *et al*., 2021). Additionally, NBR1 mediates the recruitment of TAX1BP1 to the condensates (Turco *et al*., 2021), which in turn is the main recruiter of the core autophagy protein FIP200 (Turco *et al*., 2021). A complex network of interactions then mediates the assembly and activation of the autophagy machinery including the TBK1 kinase at the p62-ubiquitin condensates (Feng *et al*, 2023; Schlütermann *et al*, 2021). Depletion of TAX1BP1 impedes the clearance of proteins aggregates in cells and mutations in this protein lead to the accumulation of ubiquitinated proteins in the brain (Sarraf *et al*, 2020).

The recruitment of TAX1BP1 to the p62-ubiquitin condensates is thus a crucial step during aggrephagy and its recruitment may trigger the switch from cargo collection in the condensates to condensate sequestration by autophagosomes. However, the mechanistic basis for the recruitment of TAX1BP1 to the condensates is still enigmatic. Through a combination of biochemical reconstitution and cell biology approaches, we reveal that TAX1BP1 is not a constitutive component of p62 condensates in cells. Dissection of the molecular basis for the interaction between TAX1BP1 and NBR1 revealed that GABARAP proteins compete with TAX1BP1 for NBR1 binding and decrease TAX1BP1 recruitment. Since ubiquitin binding appears to promote the recruitment of TAX1BP1 to p62-ubiquitin condensates, we propose a model in which TAX1BP1 acts as a ubiquitin sensor and is only recruited to p62 condensates once a certain local ubiquitin concentration threshold is exceeded.

## Results

### TAX1BP1 is not a constitutive component of p62 condensates in cells

Since TAX1BP1 is a crucial factor for the initiation of autophagosome biogenesis at the p62 condensates, we wanted to follow its dynamics in relation to p62 and NBR1 in cells. To this end, we endogenously tagged p62 with GFP, NBR1 with mScarlet and TAX1BP1 with iRFP. Comparison of the autophagic flux between the engineered and parental cell line revealed no significant difference in the stabilization of TAX1BP1, NBR1, p62 as well as the activated form of p62 phosphorylated at S349 by Bafilomycin (Baf) treatment (Fig. S1A, B). Under basal conditions p62 and NBR1 colocalized extensively when imaged by spinning disk live cell microscopy (Fig. 1A, left panel). By contrast, TAX1BP1 showed an even distribution throughout the cell and a low level of colocalization with p62 and NBR1 (Fig. 1A, left panel). However, treatment with the class III phosphatidylinositol 3-phosphate kinase (PI3KC3) inhibitor VPS34 IN1 (Bago *et al*, 2014), which blocks autophagosome biogenesis, induced foci formation for TAX1BP1 (Fig. 1A, right panel). These foci co-localized with a fraction of the p62 and NBR1 condensates (Fig. 1B, C). The finding that TAX1BP1 is not a constitutive component of the p62 condensates and that we needed to arrest autophagosome biogenesis to detect robust colocalization of TAX1BP1 with these condensates was surprising because in a fully reconstituted system in which condensate formation is induced by mixing GST-4xUB, p62 and NBR1, TAX1BP1 is recruited to almost all condensates in a NBR1 dependent manner (Fig. 1D, E)(Turco *et al*., 2021; Zaffagnini *et al*., 2018). These results suggested that there is an unknown level of regulation for the recruitment of TAX1BP1 to the p62 condensates in cells. Our data further suggested that the recruitment of TAX1BP1 may coincide with induction of autophagosome biogenesis and thus the switch from cargo collection to its sequestration by autophagosomes.

**Figure 1:**
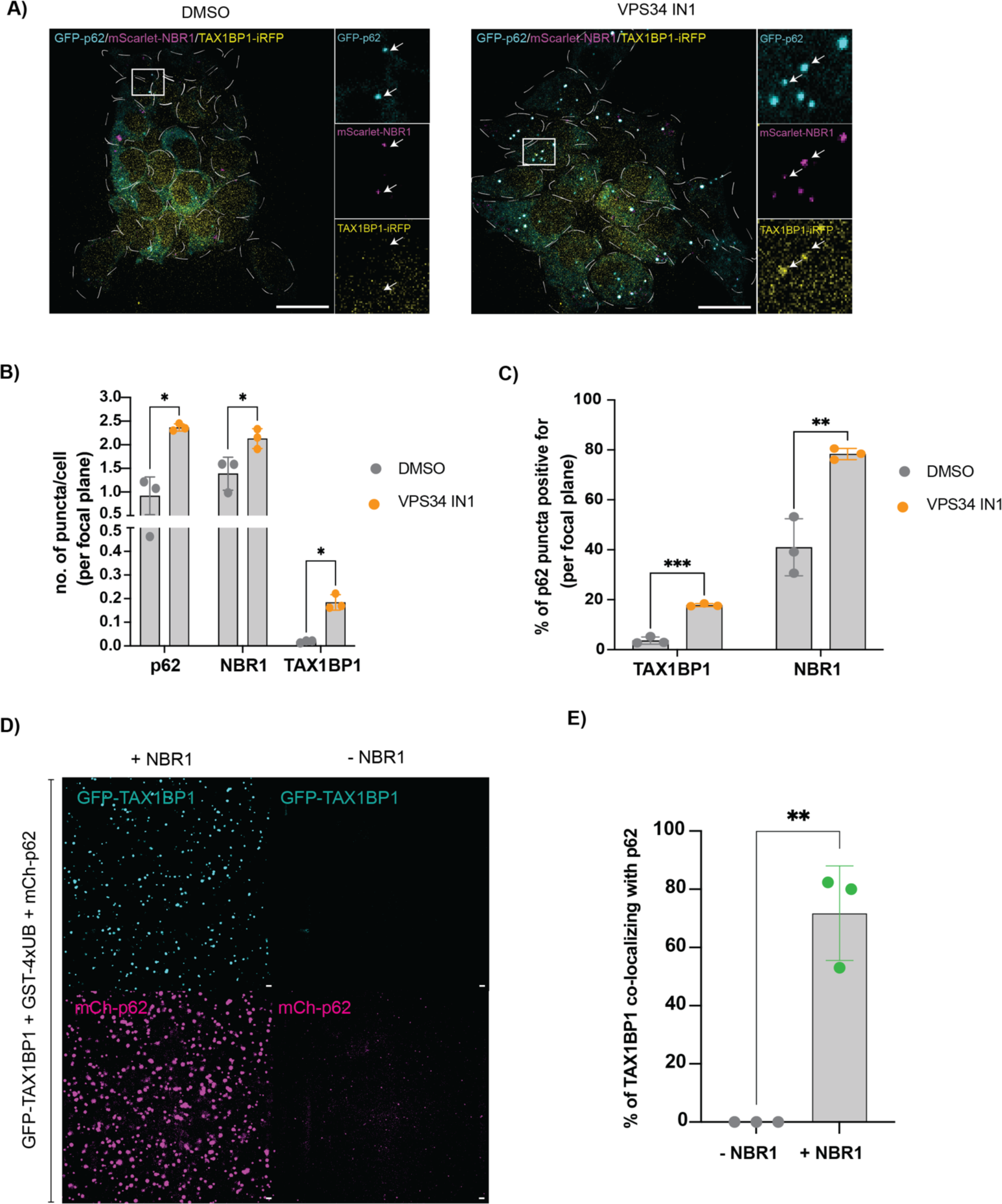
TAX1BP1 is not a constitutive component of p62 condensates in cells. A) Representative live cell images of HAP1 cells expressing endogenously tagged p62, NBR1 and TAX1BP1 untreated and upon autophagy inhibition by VPS34 IN1 (scale bar = 20 µm). B) Quantification of the number of puncta of the different cargo receptors forming with and without autophagy inhibition in A. C) Quantification of the degree of overlap between the cargo receptors in A. D) Representative image of the *in vitro* condensation assay +/− NBR1 (scale bar = 10 µm). E) Quantification of the experiment in D. Overlap between TAX1BP1 and p62 condensates at 15 min after addition of GST-4xUB is shown. Data in B, C and E show the mean +/− s.d. from three independent experiments. Unpaired t-tests were performed in B, C and E. *P<0.05, **P<0.005, ***P<0.001.

### TAX1BP1 is crucial for autophagic flux of p62 condensates

To confirm the role of TAX1BP1 as a regulator of autophagic flux of p62 condensates in our cell system, we generated a TAX1BP1 knockout (KO) in the background of a GFP-p62 expressing cell line. Employing live cell imaging, we found that the overall size and total number of p62 condensates was significantly increased in the TAX1BP1 KO cells compared to the parental cell line (Fig. 2A, B). The addition of the autophagy inhibitor VPS34 IN1 did not lead to a significant further increase in the number of p62 condensates in the TAX1BP1 KO cells, unlike in the parental cell line (Fig. 2B). Consistently, NBR1 levels were significantly increased in the TAX1BP1 KO cell line (Fig. 2C, D). Furthermore, TAX1BP1 KO cells showed a robust increase in the number of p62 condensates compared to the parental cell line upon inhibition of the proteasome by MG132 (Fig. S2A). Taken together, our data shows that autophagic flux of p62 condensates is regulated by TAX1BP1 in these cells. Next, we probed for colocalization between p62 and the autophagy machinery by immunofluorescence. This revealed a significant reduction in colocalization between p62 condensates and the autophagy core components ATG9A and FIP200 in the absence of TAX1BP1 (Fig. 2E, F)(Turco *et al*., 2021), supporting the hypothesis that TAX1BP1 is the main recruiter of the autophagy machinery.

**Figure 2:**
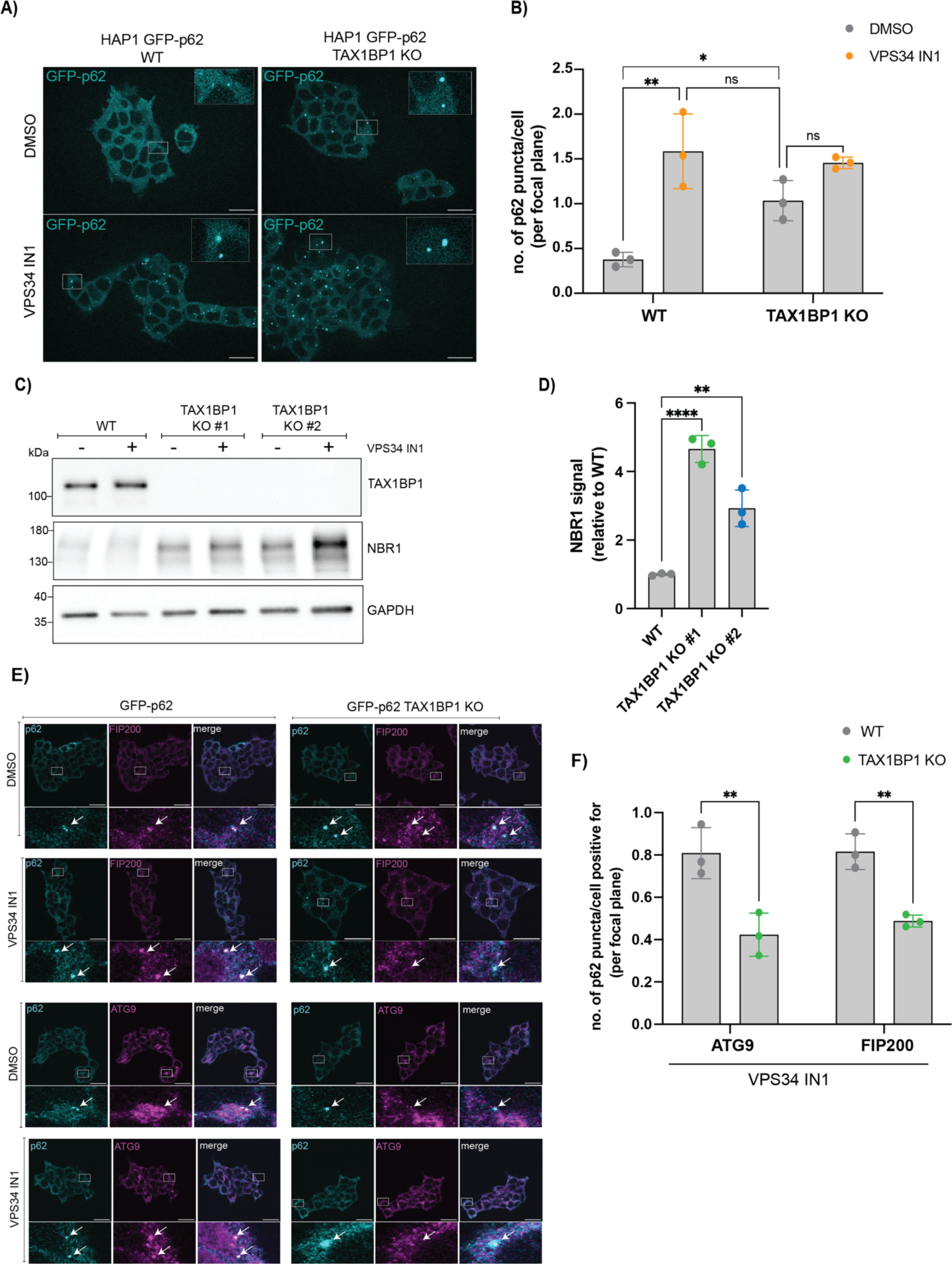
TAX1BP1 is crucial for autophagic flux of p62 condensates in cells. A) Representative live cell images of GFP-p62 expressing WT and TAX1BP1 KO cells untreated and upon VPS34 IN1 treatment (scale bar = 20 µm). B) Quantification of number of p62 puncta in A. C) Western blot of NBR1 for the WT and TAX1BP1 KO cells. D) Quantification of the western blot in C. NBR1 protein levels in control condition (DMSO) were quantified as a relative to WT. E) Representative immunofluorescence images of the WT and TAX1BP1 KO cells untreated and upon autophagy inhibition stained for FIP200 and ATG9 (scale bar = 20 µm). F) Quantification of images in E. Quantification was performed using VPS34 IN1 samples. Data in B, D and F show the mean +/− s.d. from three independent experiments. Two-way ANOVA (Analysis of variance) with Sidak’s multiple comparison test was performed in B & F. One-way ANOVA with Dunnett’s multiple comparison test was performed in D. *P<0.05, **P<0.005, ****P<0.0001, ns = not significant.

### TAX1BP1 can recruit the kinase TBK1 to p62 condensates via the adapter proteins SINTBAD/NAP1

To assess whether TAX1BP1 also mediates the recruitment of other components of the selective autophagy machinery to the condensates, we focused on TBK1. The TBK1 kinase has emerged as a major pro-autophagic factor in selective autophagy and binds to TAX1BP1 via the NAP1/SINTBAD adapters (Fu *et al*, 2018). In addition, it has recently been shown that the depletion of TAX1BP1 results in reduced TBK1 activation under basal conditions (Yamano *et al*, 2024). Indeed, we found that phosphorylation of p62 at S403, which is mediated by TBK1 (Matsumoto *et al*, 2011), was abolished in the TAX1BP1 KO cells. This was particularly evident upon inhibition of autophagy with VPS34 IN1, which resulted in the accumulation of phosphorylated p62 (Fig. 3A, B). We also observed a significant reduction in phosphorylation of the TBK1 adapter protein NAP1 (Fig. 3A, C). Thus, we speculated that TAX1BP1 might promote the recruitment of TBK1 to p62 condensates. Consistently, we found that NAP1 showed a significant reduction in colocalization with p62 condensates in the TAX1BP1 KO cells compared to the parental cell line (Fig. S3A, B). In addition, the colocalization of p62 condensates with active TBK1, phosphorylated at S172, was also reduced in the TAX1BP1 KO cells (Fig. 3D, E). These cellular data therefore suggest that TAX1BP1 is required to recruit TBK1 to p62 condensates. To test this more directly, we reconstituted these steps *in vitro*. We incubated p62, NBR1, TAX1BP1, the TBK1 adapter SINTBAD, which is structurally and functionally closely related to NAP1 (Adriaenssens *et al*, 2023) as well as TBK1. Addition of GST-4xUB induced p62-driven condensation and revealed that TAX1BP1 is both necessary and sufficient to recruit TBK1 to p62 condensates via the NAP1/SINTBAD adapters (Fig. 3F-K, S3C, S3D). Taken together our data showed that TAX1BP1 plays a major role in mediating TBK1 recruitment to p62 condensates and thus potentially promotes the local phosphorylation of p62 and other components in these structures (Fig. 3K).

**Figure 3:**
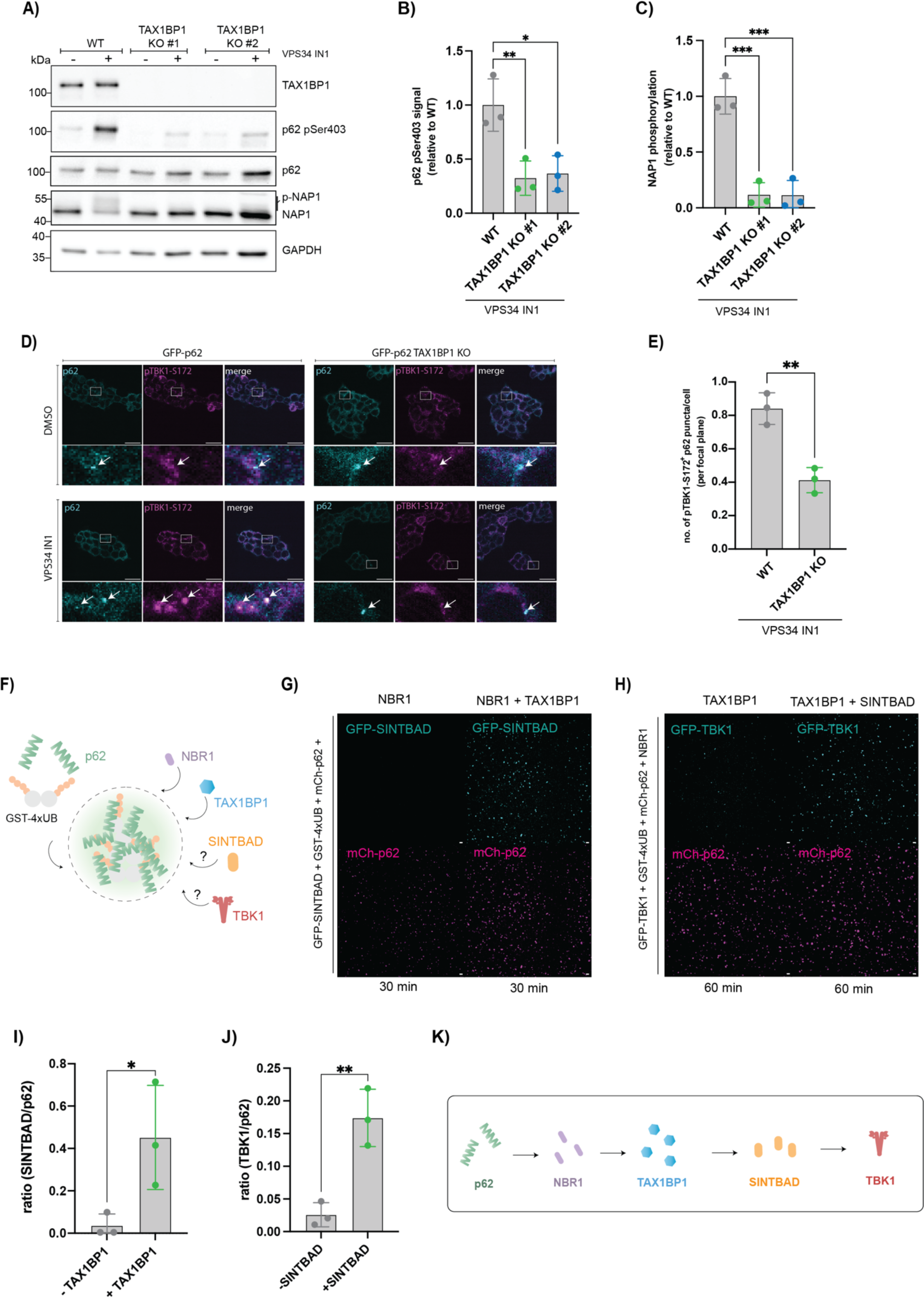
TAX1BP1 recruits the TBK1 kinase to p62 condensates via the adapter proteins SINTBAD/NAP1. A) Western blots for p62, phosphorylated p62 and the adapter protein NAP1. B) Quantification of western blot in A. Protein levels of phosphorylated p62 upon autophagy inhibition were quantified as a relative to WT. C) Quantification of western blot in A. Protein levels of phosphorylated NAP1 upon autophagy inhibition were quantified as a relative to WT. D) Representative immunofluorescence images of cells stained for p62 and TBK1 Ser172 untreated and upon autophagy inhibition (scale bar = 20 µm). E) Quantification of the number of p62 puncta positive for TBK1 Ser172 upon autophagy inhibition in D. F) Scheme of the *in vitro* condensation assay. G) Representative image of the *in vitro* reconstitution of GFP-SINTBAD recruitment to p62 condensates +/− TAX1BP1 (scale bar = 10 µm; loading control in Fig. S3C). H) Representative image of the *in vitro* reconstitution of GFP-TBK1 recruitment to p62 condensates +/− SINTBAD (scale bar = 10 µm; loading control in Fig. S3D). I) Quantification of G. Ratio between number of SINTBAD and p62 condensates is shown at 30 minutes after addition of GST-4xUB. J) Quantification of H. Ratio between number of TBK1 and p62 condensates is shown at 60 minutes after addition of GST-4xUB. K) Schematic summary of the results from the condensation experiments. Data in B, C, E, I and J show the mean +/− s.d. from three independent experiments. One-way ANOVA with Dunnett’s multiple comparison test was performed in B and C. Unpaired t-test was performed in E, I and J. *P<0.05, **P<0.005, ***P<0.001, ****P<0.0001.

### TAX1BP1 binds to the LIR motif of NBR1 via its N-Domain *in vitro*

How TAX1BP1 is recruited to the p62 condensates is not well understood. While it was shown that the interaction with NBR1 requires a central part of TAX1BP1, referred to as N-domain (aa420-506) (Ohnstad *et al*, 2020), the mechanistic basis of this interaction remains unexplored. We previously showed that the FW domain of NBR1 is sufficient for the interaction with TAX1BP1 (Turco *et al*., 2021). However, the FW domain is not essential because its deletion did not abolish the binding to TAX1BP1. We therefore set out to systematically define the interaction between TAX1BP1 and NBR1 with the aim to reveal the basis of a potential regulatory mechanism. We started by testing if the TAX1BP1 N-domain is not only required but also sufficient for the interaction with NBR1. Employing a microscopy-based protein-protein interaction assay we found that the N-domain is indeed sufficient to recruit NBR1 (Fig. 4A, B, S4A). Next, we determined the corresponding binding region in NBR1. Multiple attempts to define this interaction site using AlphaFold 2 gave inconclusive results. We therefore used a peptide array where we spotted NBR1 peptides on a membrane. Each peptide consisted of 12 amino acids with two residues overlap between peptides. We then probed for interacting peptides with recombinant TAX1BP1. We identified two regions capable of binding to TAX1BP1: parts of the FW domain, previously implicated to bind TAX1BP1 weakly (Turco *et al*., 2021) and the LC3-interacting region (LIR) 1 (Kirkin *et al*., 2009) (designated as binding region 1 and 2, respectively; aa347 – aa365 and aa731 - aa749) (Fig.4C). Next, we tested if the deletion of the identified binding regions would abrogate the binding between NBR1 and TAX1BP1. Transfection of cells with corresponding deletions of NBR1 and subsequent pulldowns from the cell lysates using GST-TAX1BP1 immobilized on bead, demonstrated that binding region 2, encompassing the LIR1 of NBR1, is required for the interaction while binding region 1 is not (Fig. 4D). We also confirmed that a small fragment of NBR1 (referred to as CC2-LIR1; aa685 – aa757), which contains the binding region 2, alone is sufficient for the interaction with TAX1BP1 *in vitro* (Fig. S4B, C). To obtain further insights into the interaction of the NBR1 LIR1 region with the N-domain of TAX1BP1, we performed AlphaFold 2 modeling with sequences of TAX1BP1 and NBR1 encompassing the identified binding sites (Fig. 4E, S4D)(Jumper *et al*, 2021). Based on the highest-ranking model, we generated point mutants for both TAX1BP1 and NBR1 and assessed their impact on the interaction of the two proteins. TAX1BP1 point mutants were validated in a microscopy-based interaction assay (Fig. 4F, S4E). NBR1 point mutants were validated performing a pulldown with cell lysates (Fig. 4H). Residues Y449 and L442 in TAX1BP1 and F740 of NBR1 were identified to be essential for the interaction between the two proteins, consistent with our AF2 model (Fig. 4F, G, H, S4E). Given the proximity of residue F740 to the LIR1 motif, we tested whether it is also required for binding to GABARAP/LC3 proteins. To this end, we performed a pulldown with GST-LC3B which resulted in only a mild reduction in LC3B binding (Fig. S4F).

**Figure 4:**
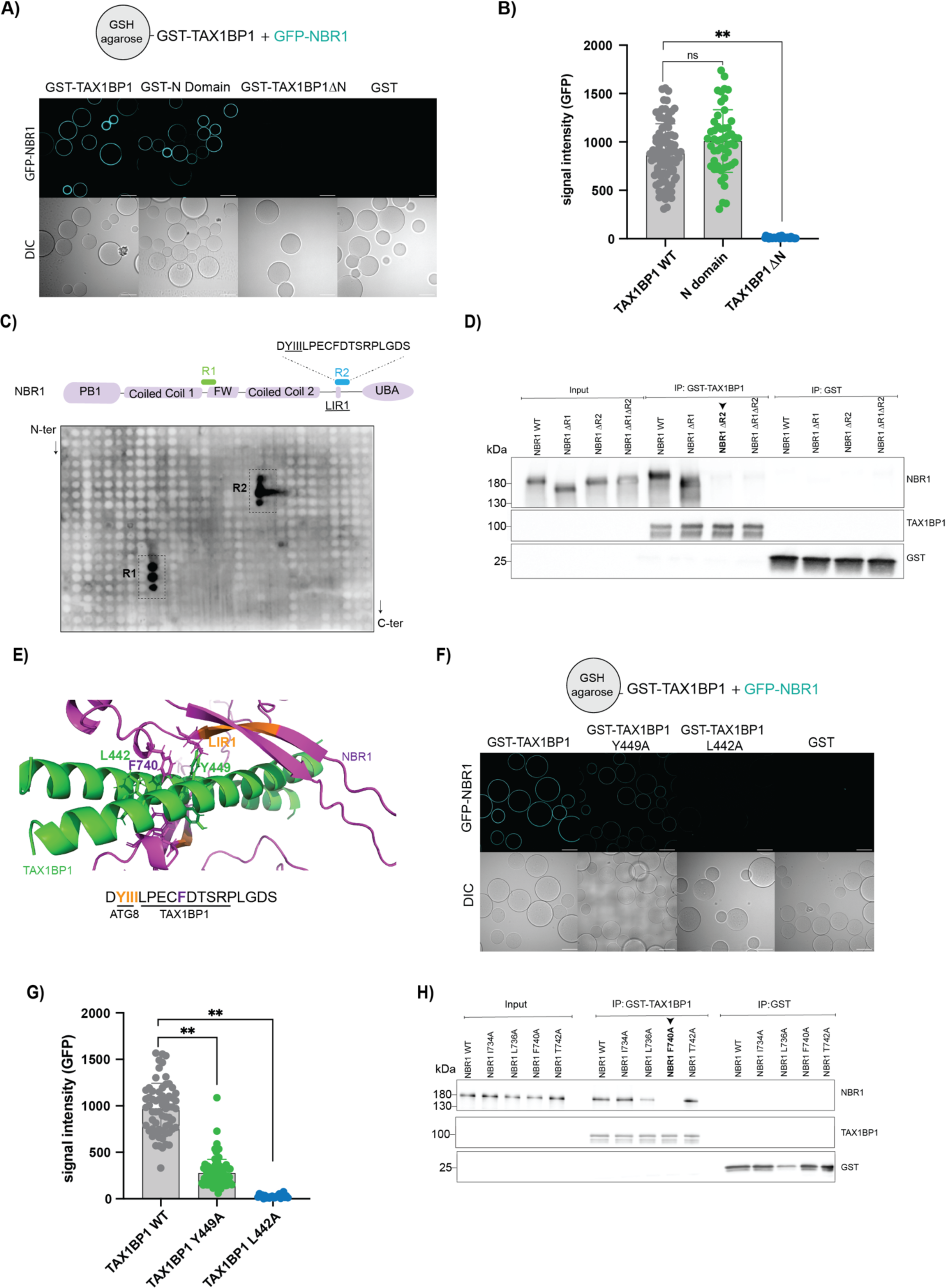
TAX1BP1 binds to the LIR motif of NBR1 via its N-domain. A) Representative images of a microscopy-based protein-protein interaction assay (scale bar = 80 µm; loading control in Fig. S4A). B) Quantification of signal intensity of GFP-NBR1 in A. C) Western blot of NBR1 peptide array after incubation with recombinant TAX1BP1 protein. D) Western blot of pulldown with cells expressing NBR1 truncations. E) AlphaFold 2 model of the interaction between the TAX1BP1 N-domain and the binding region 2 including the LIR1 of NBR1 (pLDDT 44.3 and ptmscore 0.333 / iptm 0.387). The region of NBR1 encompassing aa684 – aa966 was modeled together with residues aa430 – aa468 of TAX1BP1 in a dimeric form using AlphaFold 2. F) Representative images of a microscopy-based protein-protein interaction assay using TAX1BP1 point mutants (scale bar = 80 µm; loading control in Fig. S4E). G) Quantification of signal intensity of GFP-NBR1 in F. H) Western blot of pulldown with cells expressing NBR1 point mutants and GST-TAX1BP1 immobilized on beads. Data in B and G show the mean +/− s.d. from three independent experiments. Each data point in B and G represent the mean signal intensity for an individual bead. Nested one-way ANOVA with Dunnett’s multiple comparison test was performed in B and G. **P<0.005, ns = not significant.

### GABARAP competes with TAX1BP1 for NBR1 binding

Based on the discovery that the LIR1 region of NBR1 is essential for the binding to TAX1BP1, we speculated that the interaction of NBR1 with TAX1BP1 and ATG8 proteins such as GABARAP might be mutually exclusive. To test this, we co-incubated GFP-NBR1 coated beads simultaneously with mCherry-TAX1BP1 and increasing amounts of GABARAP. Binding of TAX1BP1 to NBR1 was gradually reduced by the addition of increasing concentrations of GABARAP (Fig. 5A, B, S5A). The addition of GABARAP also outcompeted an already established TAX1BP1-NBR1 interaction *in vitro* (Fig 5C, D). Next, we asked if GABARAP would also reduce the recruitment of TAX1BP1 to p62 condensates. Indeed, the presence of GABARAP notably reduced the fraction of TAX1BP1 positive p62 condensates in our condensation assay (Fig. 5E, F, S5B).

**Figure 5:**
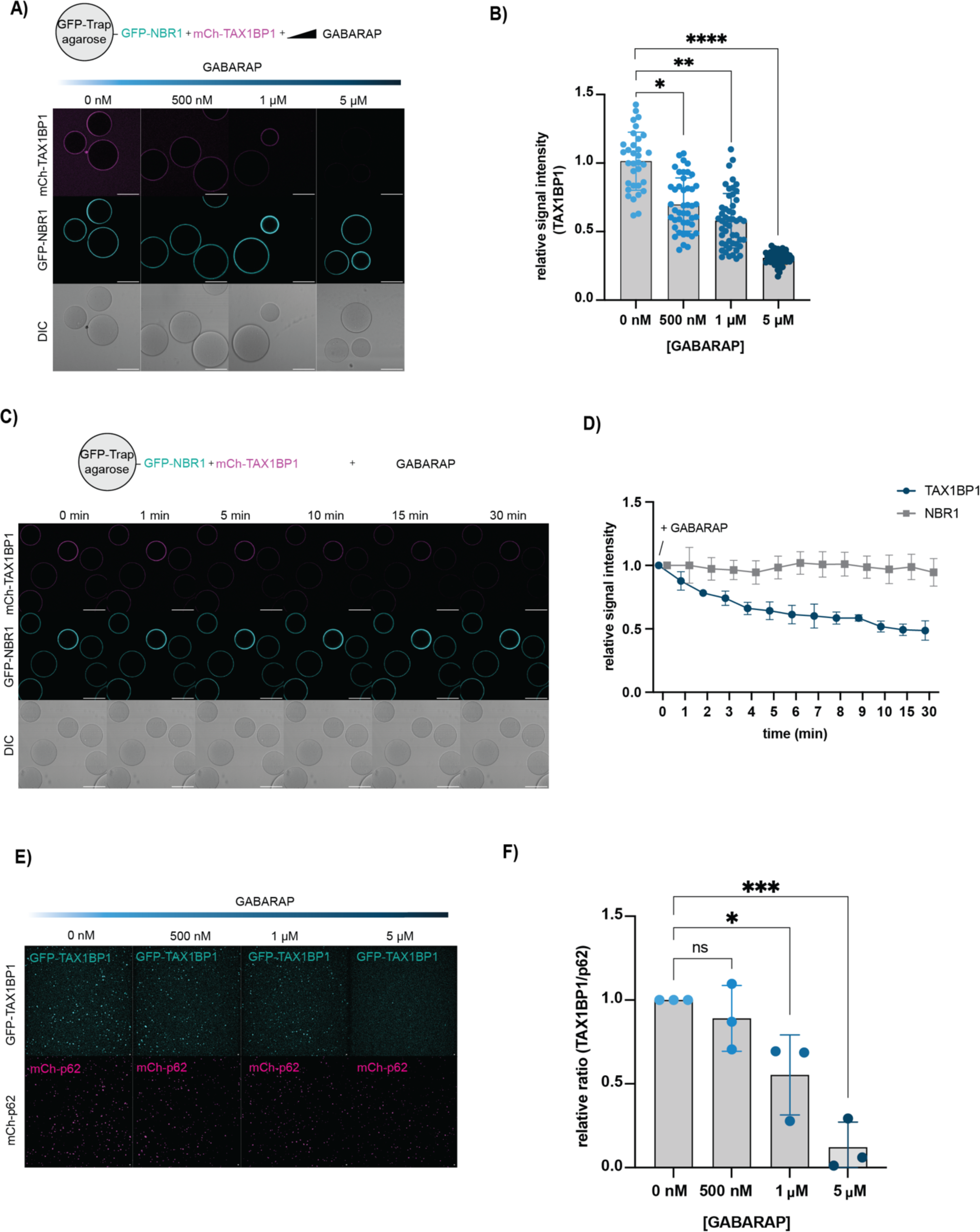
GABARAP competes with TAX1BP1 for binding to NBR1. A) Representative images of a microscopy-based protein-protein interaction assay adding increasing amounts of GABARAP protein (scale bar = 80 µm; loading control in Fig. S5A). B) Quantification of signal intensity of mCh-TAX1BP1 in A as a relative to the ctrl (0 nM GABARAP). C) Representative images of a microscopy-based protein-protein interaction assay adding 5 µM GABARAP protein to GFP-NBR1 coupled beads pre-incubated with mCh-TAX1BP1 to monitor the loss of TAX1BP1 signal over time (scale bar = 80 µm). D) Quantification of mCh-TAX1BP1 signal in C as a relative to timepoint 0 (Addition of GABARAP). E) Representative images of the *in vitro* condensation assay adding increasing amounts of GABARAP protein (scale bar = 10 µm; loading control in Fig. S5B) F) Quantification of F. Ratio between number of p62 and TAX1BP1 condensates was plotted as a relative to ctrl (0 nM GABARAP). Data in B, D and F show the mean +/− s.d. from three independent experiments. Each data point in B and F represent the mean signal intensity for an individual bead. One-way ANOVA with Dunnett’s multiple comparison test was performed in B and F. *P<0.05, **P<0.005, ***P<0.001, ****P<0.0001, ns = not significant

Thus, our findings delineate the interaction between NBR1 and TAX1BP1, revealing that it involves the LIR1 region of NBR1 and that the interaction can be outcompeted by high concentrations of LC3/GABARAP proteins.

### The TAX1BP1 – NBR1 interaction is required for the autophagic flux of p62 condensates

To test the relevance of the interaction between TAX1BP1 and NBR1 for the autophagic degradation of p62 condensates in cells, we re-introduced fluorescently tagged wild-type (WT), ΔN (lacking the N-domain) and L442A TAX1BP1 into the TAX1BP1 KO cell line by stable transfection (Fig. S6A). After doxycycline-induced expression, we observed that in contrast to TAX1BP1 WT (R-WT) and consistent with our *in vitro* data, TAX1BP1 ΔN and L442A (R-ΔN, R-L442A) failed to colocalize with p62 condensates upon treatment with VPS34 IN1 (Fig. 6A, B). Intriguingly, even upon overexpression, TAX1BP1 did not co-localize with p62 puncta and the number of TAX1BP1 foci upon VPS34 IN1 remained lower than for p62 (Fig. 6A, S6B). Furthermore, re-expression of TAX1BP1 WT rescued the phenotype of the TAX1BP1 KO reducing the number of p62 condensates, while the mutants failed to do so (Fig. 6C). Additionally, the TAX1BP1 mutants were unable to restore TBK1-mediated phosphorylation of p62 and the adapter protein NAP1 (Fig. 6D-F). Taken together, the interaction between TAX1BP1 and NBR1 is crucial for TBK1 recruitment and to promote autophagy flux of p62 condensates in cells.

**Figure 6:**
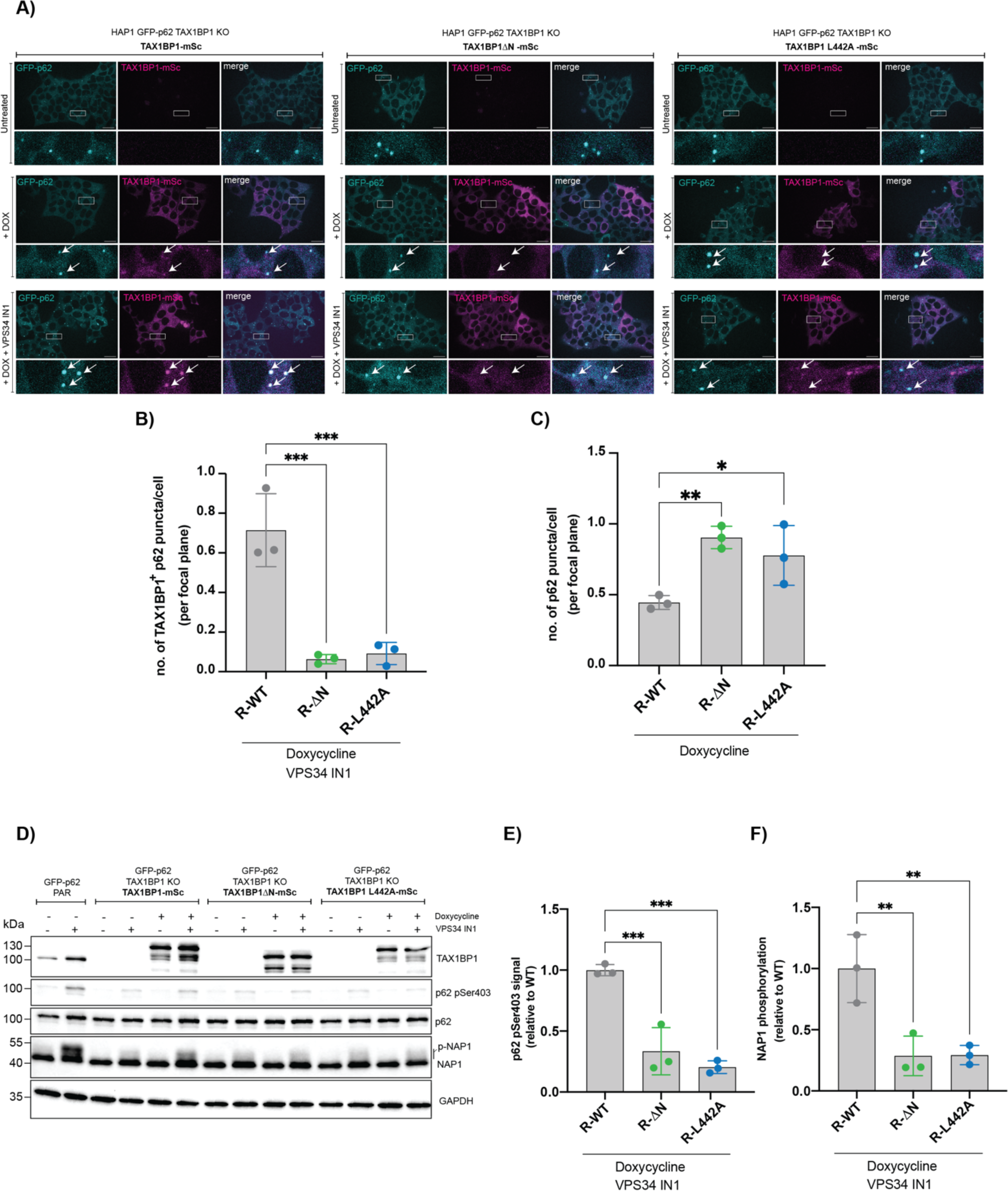
The TAX1BP1-NBR1 interaction is required for the autophagic flux of p62 condensates in cells. A) Representative live cell images of cells untreated, upon doxycycline treatment to induce expression of TAX1BP1 and upon autophagy inhibition (scale bar = 20 µm). B) Quantification of the number of p62 puncta positive for TAX1BP1 upon addition of doxycycline and VPS34 IN1 from A. C) Quantification of total number of p62 puncta upon addition of doxycycline from A. D) Western blot of cells expressing TAX1BP1 WT, ΔN or L442A upon doxycycline treatment and autophagy inhibition. E) Quantification of western blot in D. Phosphorylation levels of p62 were quantified as a relative to R-WT. F) Quantification of western blot in D. Phosphorylation levels of NAP1 were quantified as a relative to R-WT. H) Western blot of a pulldown with cells expressing NBR1 point mutants. Data in B, C, E and F show the mean +/− s.d. from three independent experiments. One-way ANOVA with Dunnett’s multiple comparison test was performed in B, C, E and F. *P<0.05, **P<0.005, ***P<0.001.

### Ubiquitin levels can regulate TAX1BP1 recruitment to p62 condensates

Based on the insights we obtained regarding the TAX1BP1 – NBR1 interaction, we aimed to understand how the TAX1BP1 recruitment and by implication the initiation autophagosome biogenesis is regulated. p62 condensates harbor two known binding cues for TAX1BP1: ubiquitin, which is bound by TAX1BP1 via its C-terminal ZnF domain (S7A-C)(Ferrari *et al*, in press; Sarraf *et al*., 2020; Tumbarello *et al*, 2015) and the region around the LIR1 motif of NBR1, which we identified above. To identify the respective contribution of each individual binding cue for the recruitment of TAX1BP1 to p62 condensates, we reconstituted this step *in vitro*. Deletion of the N-domain completely abolished the recruitment of TAX1BP1 to p62 condensates, while the N-domain alone was still recruited, albeit to a lower level (Fig. 7A, B, S7D). Conversely, deletion of the ZnF domain resulted in a loss of ubiquitin binding (Fig. S7A-C) and a decreased TAX1BP1 recruitment to p62 condensates *in vitro* (Fig. 7B), consistent with recent findings (Ferrari *et al*., in press). The levels of recruitment of TAX1BP1ΔZnF and of the isolated N-domain were similar, suggesting that the residual recruitment of TAX1BP1ΔZnF is mediated by the N-domain (Fig. 7A, B, S7D). To further corroborate our *in vitro* results, we re-introduced TAX1BP1ΔZnF (labelled as R-ΔZnF) into our TAX1BP1 KO cell line in a doxycycline-inducible manner. In line with our *in vitro* result, we observed a mild reduction in TAX1BP1 puncta overlapping with p62 and weaker NAP1 phosphorylation levels in comparison to the TAX1BP1 WT (Fig. 7C-F). These results suggest that under the conditions tested, the N-domain-mediated interaction with NBR1 is essential for TAX1BP1 recruitment, while the ZnF mediated recruitment via ubiquitin merely enhances the recruitment or stabilizes the interaction.

**Figure 7:**
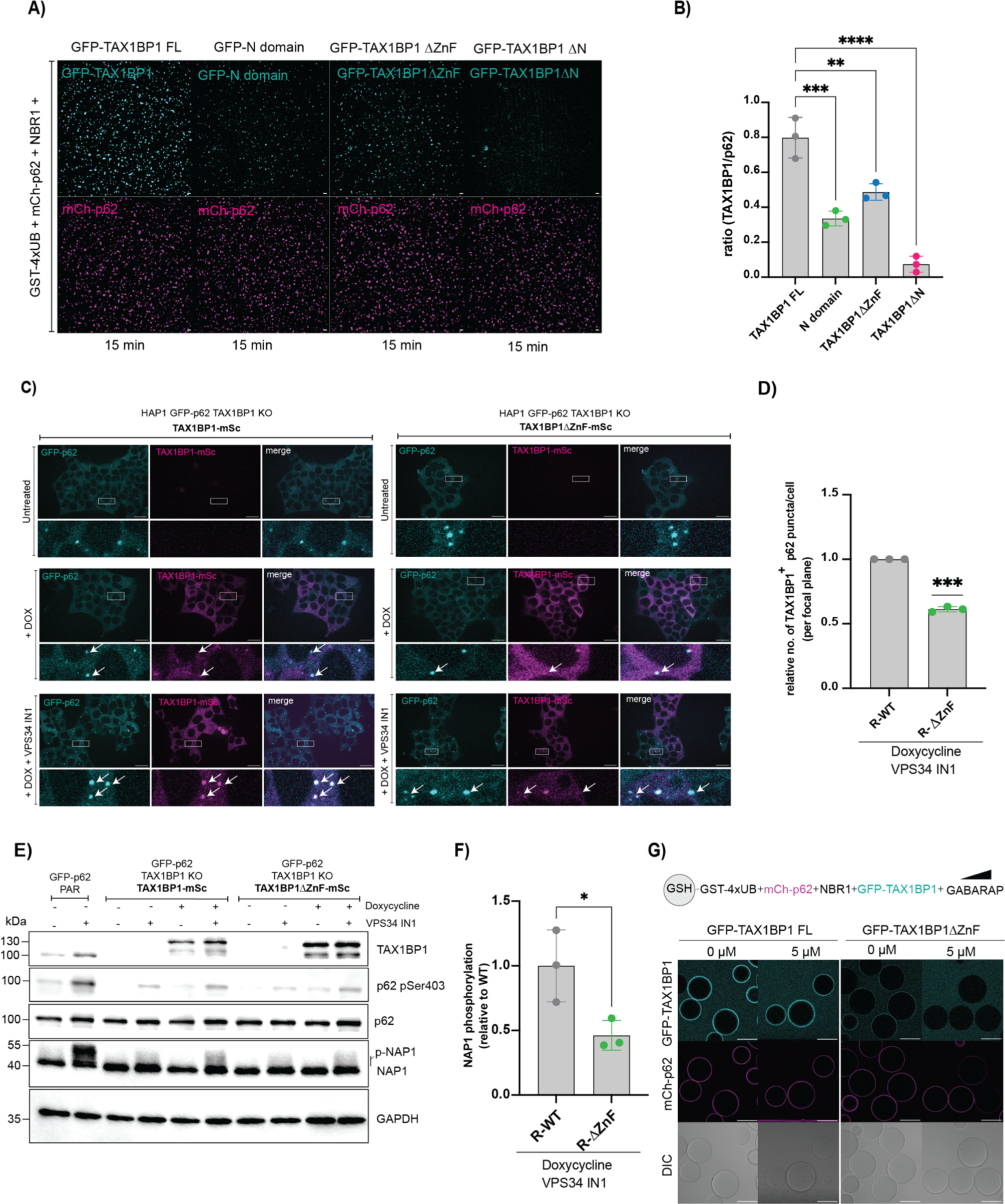
Ubiquitin interaction stabilizes TAX1BP1 at p62 condensates *in vitro* and in cells. A) Representative images of the *in vitro* condensation assay comparing the recruitment of TAX1BP1 full length (FL) vs ΔN vs ΔZnF (scale bar = 10 µm; loading control in Fig. S7D). B) Quantification of A. Ratio of number of p62 condensates and GFP-TAX1BP1 (FL/N domain/TAX1BP1ΔZnF/TAX1BP1ΔN) are plotted and compared. C) Representative live cell images of cells untreated, upon doxycycline treatment to induce expression of TAX1BP1 and upon autophagy inhibition (scale bar = 20 µm). D) Quantification of the number of p62 puncta positive for TAX1BP1 upon addition of doxycycline and VPS34 IN1 as a relative to R-WT from C. E) Western blot of cells expressing TAX1BP1 WT or ΔZnF upon doxycycline treatment and autophagy inhibition with VPS34 IN1. F) Quantification of western blot in E. NAP1 phosphorylation levels were quantified as a relative to R-WT. G) Representative images of microscopy-based interaction assay to compare TAX1BP1 recruitment to cargo mimetic beads in the presence of GABARAP with and without ubiquitin binding (scale bar = 80 µm; loading control in Fig. S7E). Data in B, D and F show the mean +/− s.d. from three independent experiments. One-way ANOVA with Dunnett’s multiple comparison test was performed in B. One-sample t-test and Wilcoxon-test was performed in D. Unpaired t-test was performed in F. *P<0.05, **P<0.005, ***P<0.001, ****P<0.0001.

Since we found above that the binding of TAX1BP1 to NBR1 is weakened in the presence of GABARAP (Fig. 5), we tested whether the ZnF-mediated interaction with ubiquitin aids in overcoming the competition with GABARAP for efficient TAX1BP1 recruitment. We performed reconstitution experiments using cargo-mimetic beads coated with GST-4xUB and compared the recruitment of GFP-TAX1BP1 full-length (FL) and ΔZnF to the beads in the presence of p62, NBR1 and increasing amounts of GABARAP. To ensure that ubiquitin remains a limiting factor and that TAX1BP1 recruitment depends also on NBR1 (as observed in cells (Fig. 6A-C)), we employed low amounts of ubiquitin and pre-coated the beads with mCherry-p62 and NBR1. Indeed, under the conditions tested, the full-length TAX1BP1 protein was robustly recruited to the beads even in the presence of high concentrations of GABARAP while TAX1BP1ΔZnF was barely detectable (Fig. 7G, S7E). This suggests that due to their competition with TAX1BP1 for NBR1 binding, the GABARAP/LC3 proteins render the ubiquitin interaction more important for TAX1BP1 recruitment. Hence, we propose that TAX1BP1 is recruited to p62 condensates once a certain local ubiquitin threshold is surpassed, to trigger the switch from cargo collection to autophagosome biogenesis when the cargo load reaches a sufficient level (Fig. 8).

**Figure 8:**
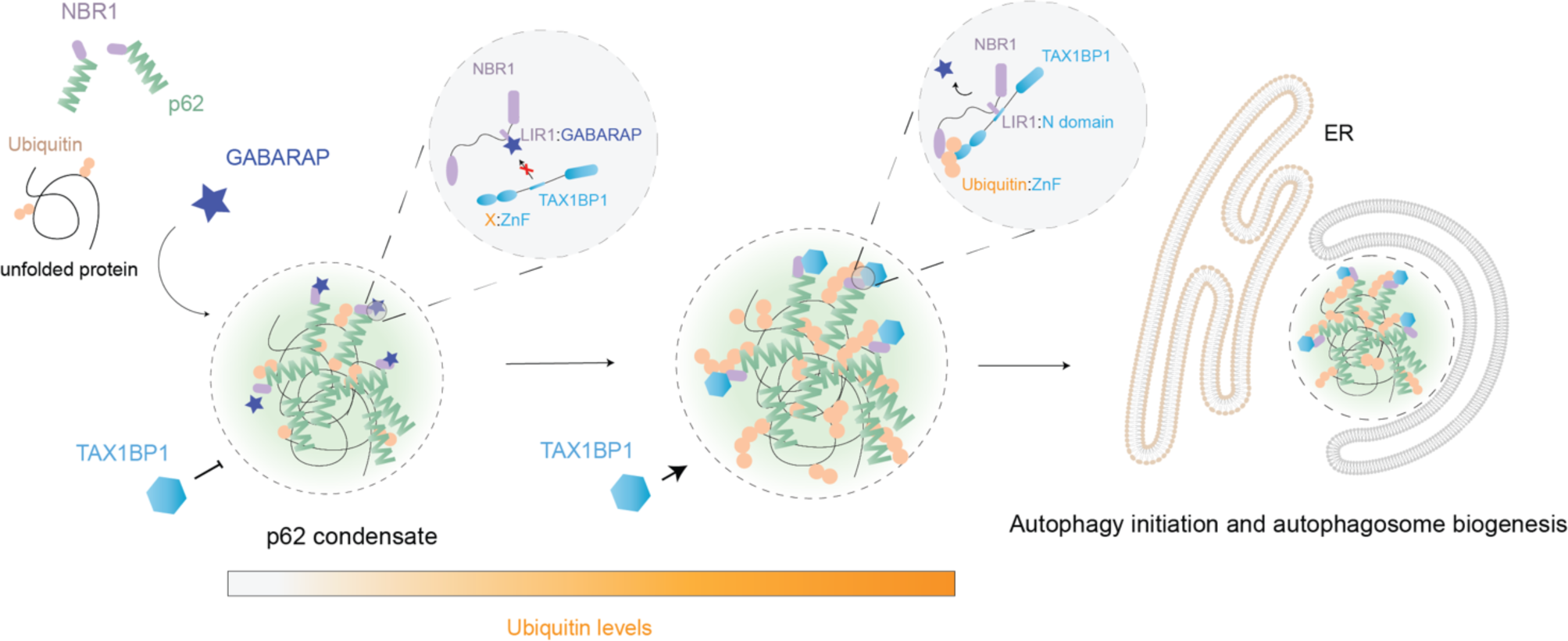
Model for the ubiquitin load dependent, TAX1BP1-mediated switch from cargo collection to its sequestration in aggrephagy.

## Discussion

Selective autophagy is a major pathway for the degradation of cellular structures that are out of reach of other degradation systems. For example, it can mediate the degradation of complex organelles such as mitochondria, the endoplasmic reticulum, peroxisomes and even pathogens (Vargas *et al*, 2023). These cargoes have in common that they represent large structures to which the autophagy machinery can be recruited by cargo receptors to induce autophagosome biogenesis (Ravenhill *et al*, 2019; Vargas *et al*, 2019). In this respect aggrephagy is distinct from other selective autophagy pathways, as the cargo material must first be collected into larger condensates before autophagy is triggered. The main collector of the cargo material in aggrephagy is p62 (Bjorkoy *et al*., 2005; Ciuffa *et al*., 2015; Kageyama *et al*., 2021; Sarraf *et al*., 2020; Sun *et al*., 2018; Turco *et al*., 2021; Zaffagnini *et al*., 2018). Various cellular materials have been shown to be accumulated in the p62 condensates (Zellner *et al*, 2021) including translation initiation factors (Danieli *et al*, 2023), aggregation-prone proteins (Bjorkoy *et al*., 2005), vault complexes (Kurusu *et al*, 2023) and negative regulators of NRF2 signaling (Kageyama *et al*., 2021). A prerequisite for the sequestration of most of these materials appears to be their ubiquitination (Bjorkoy *et al*., 2005; Sun *et al*., 2018; Zaffagnini *et al*., 2018). The requirement for cargo collection prior to its sequestration may explain why in aggrephagy at least three cargo receptors cooperate to drive this pathway. Among the three cargo receptors TAX1BP1 is the main recruiter and activator of the autophagy machinery (Sarraf *et al*., 2020; Turco *et al*., 2021), although p62 and NBR1 can also bind FIP200 (Turco *et al*., 2021; Turco *et al*, 2019). Importantly, for Alzheimer’s brain derived Tau fibrils, the recruitment of TAX1BP1 is compromised (Ferrari *et al*., in press).

Given the importance of TAX1BP1 recruitment to cargo material, we set out to understand how its activity can be regulated in cells. We discovered that, in contrast to the *in vitro* reconstituted p62/NBR1 condensates, TAX1BP1 is not a constitutive component of these structures in cells. This finding suggests that TAX1BP1 recruitment is a major regulatory mechanism. Furthermore, we found that the inhibition of autophagosome biogenesis after its induction using the PI3KC3C1 inhibitor IN1 was required to robustly detect TAX1BP1 at the condensates. Together with the result that deletion of TAX1BP1 results in the accumulation of p62 condensates (Sarraf *et al*., 2020; Turco *et al*., 2021) this suggests that its recruitment coincides with the induction of autophagosomes biogenesis. Indeed, *in vitro* TAX1BP1 is both necessary and sufficient to recruit the TBK1 kinase (Fig. 3F-K), a key factor in selective autophagy as well as FIP200 (Adriaenssens *et al*., 2023; Matsumoto *et al*., 2011; Nguyen *et al*, 2023; Richter *et al*, 2016; Turco *et al*., 2021).

We therefore determined the molecular basis for the recruitment of TAX1BP1 to the p62/NBR1 condensates. It was previously shown that a region named N-domain encompassing amino acids 420-506 of TAX1BP1 is required for its interaction with NBR1 (Ohnstad *et al*., 2020). Furthermore, NBR1 is required for the recruitment of TAX1BP1 to p62 condensates (Turco *et al*., 2021). We found that the N-domain is indeed sufficient for the binding to NBR1. To our surprise we found that the main binding site for the N-domain of TAX1BP1 in NBR1 is a region encompassing its LIR1 motif, which binds GABARAP/LC3 proteins (Kirkin *et al*., 2009; Pankiv *et al*, 2007). The FW domain of NBR1, which we previously showed to weakly bind to TAX1BP1 (Turco *et al*., 2021) and which also overlaps with binding region 2 (Fig. 4C) did not detectably contribute to the interaction with TAX1BP1 in our experiments. The information that the region around the LIR1 of NBR1 is necessary and sufficient for the binding to TAX1BP1 allowed us to model the interaction using AlphaFold 2. While the details of this model will have to be confirmed by structural studies, we could corroborate it through point mutations in both TAX1BP1 and NBR1. Notably, in parallel to us another study identified the same LIR1 region of NBR1 as the interaction site for TAX1BP1 (North *et al*, 2024).

The proximity of the TAX1BP1 and GABARAP/LC3 binding sites in NBR1 render the interaction of NBR1 with the two binding partners mutually exclusive. As a result, the interaction of TAX1BP1 and NBR1 is likely to be weakened in cells, where the concentration of the LC3/GABARAP proteins is considerably higher than that of TAX1BP1 and NBR1 (Cho *et al*, 2022). This also implies that in cells, TAX1BP1 recruitment might be more dependent on the second binding cue i.e. ubiquitin binding. In fact, NBR1 and ubiquitin binding by TAX1BP1 are both required for full recruitment *in vitro* as well as in cells and likely also in the brain (Sarraf *et al*., 2020). We found in a reconstituted system that the presence of GABARAP renders ubiquitin binding by TAX1BP1 essential for its recruitment to cargo mimetic beads (Fig. 7G). We therefore hypothesize that the ubiquitin dependence for its recruitment offers TAX1BP1 the opportunity to sample the ubiquitin load and by implication the cargo load of the p62/NBR1 condensates. Once this load increases above a certain threshold, TAX1BP1 is recruited and further recruits the autophagy machinery including TBK1 via the NAP1 and SINTBAD adapters as well as FIP200 (Turco *et al*., 2021) to trigger the crucial switch from cargo collection to autophagosome biogenesis (Fig. 8).

It is curious and currently unexplained why the TAX1BP1 – NBR1 interaction has evolved to be sensitive to the concentrations of LC3/GABARAP proteins rather than being of lower affinity, which would also allow TAX1BP1 to sense the ubiquitin load. GABARAP/LC3 proteins are themselves consumed by autophagy but also locally concentrated on the nascent autophagosomal membranes. Thus, their overall depletion may globally enhance the NBR1-TAXBP1 interactions while their clustering on the membrane may locally dissociate it, potentially aiding membrane bending and dynamics. Future studies will have to elucidate the molecular gymnastics of these interaction modalities in space and time during aggrephagy.

## Materials & Methods

### Cell Culture

All mammalian cell lines were cultured at 37°C in a humidified 5% CO2 atmosphere. HeLa PentaKO cells were obtained from Michael Lazarou (Lazarou *et al*, 2015). HAP1 cells were originally purchased from Horizon Discovery. HAP1 GFP-p62 and GFP-p62 mSc-NBR1 were previously generated and used as parental cell lines in this study (Turco *et al*., 2021; Zaffagnini *et al*., 2018). HeLa and HEK293T cells were grown in Dulbecco Modified Eagle Medium (DMEM, Thermo Fisher) supplemented with 10% (v/v) Fetal Bovine Serum (FBS, Thermo Fisher), 25 mM HEPES (Thermo Fisher), 1% (v/v) non-essential amino (NEAA, 11140050, Thermo Fisher), and 1% (v/v) Penicillin-Streptomycin (Thermo Fisher). HAP1 cells were cultured in Iscove’s Modified Dulbecco’s Medium (IMDM, Thermo Fisher), supplemented with 10% (v/v) Fetal Bovine Serum (FBS, Thermo Fisher) and 1% (v/v) Penicillin-Streptomycin (15140122, Thermo Fisher).

Sf9 insect cells were cultured as a shaking liquid culture at 27°C in insect culture medium (ESF 921 Expression System LLC) supplemented with 1% Penicillin-Streptomycin (Thermo Fisher).

### Reagents & Cell Treatments

The following chemical compounds have been used in this study: Bafilomycin A1 at 400 nM for 3h (sc-201550, Santa Cruz Biotech), Doxycycline at 50 ng/ml for 16h (Sigma), DMSO (D2438, Sigma), Puromycin at 5 µg/ml (Thermo Fisher), VPS34 IN1 at 2 µM for 3h (APE-B6179, ApexBio).

### Cloning

The sequences for all the inserts were generated by amplifying existing plasmids or through gene synthesis (GenScript). Plasmids were generated either by Gibson assembly or gene synthesis. For Gibson assembly, the vector and inserts were generated by restriction digestion or PCR. DNA was cleaned up with Promega Wizard SV gel and PCR Cleanup System (Promega). The inserts and the backbone were mixed in a 3:1 ratio and incubated with the 2x NEBuilder HiFi DNA assembly enzyme mix (New England Biolabs) for 1h at 50°C. Competent DH5a *E.coli* were transformed with the Gibson reaction and incubated overnight at 37°C. The next day single colonies were picked, grown overnight, and plasmids were isolated using the GeneJet Plasmid Miniprep Kit (Thermo Fisher). For sequence verification, the isolated plasmids were sent for Sanger sequencing (MicroSynth AG).

### CRISPR/Cas9 genome editing

sgRNAs for generating HAP1 GFP-p62 mSc-NBR1 TAX1BP1-iRFP cell line were designed by CRISPOR (http://crispor.tefor.net). Both sgRNAs were cloned into an all-in-one vector encoding Cas9 D10A nickase (Addgene) with puromycin as a selection marker. The plasmid bearing the repair template was generated by gene synthesis (GenScript). All-in-one vector and repair template were transfected into GFP-p62 mSc-NBR1 (previously validated in (Turco *et al*., 2021)) using Fugene 6.

After selection with puromycin, remaining cells were single-cell sorted and validated by PCR, western blot and Sanger sequencing. Selected clone was also validated for protein levels and autophagy flux.

The same sgRNAs for generating the TAX1BP1 KO cell line has been used as previously reported (Sarraf *et al*., 2020) and cloned into pSpCas9(BB)-2A-Puro vector backbone. The plasmids were transfected into HAP1 GFP-p62 cells (previously described in (Zaffagnini *et al*., 2018)). After selection with puromycin, cells were single-cell sorted. Individual clones were validated using PCR, western blots and Sanger sequencing.

### Generation of stable cell line

For stable cell line generation, lentiviral transductions were performed. HEK293T cells were transfected with VSV-G, GAG-POL and pInducer lentiviral plasmids carrying the transgene using Lipofectamine 3000 (Thermo Fisher). After 24h, the medium of the transfected cells was exchanged to the appropriate medium for the target cells. HAP1 GFP-p62 TAX1BP1 KO cells were seeded on a 6 well plate at 400k cells/well. 24h later the medium containing the lentivirus was collected and filtered with a 0.45 µm filter. 8mg/ml polybrene was added to the viral supernatant. The target cells were incubated for 24h with the virus containing medium. The cells were washed with PBS on the next day and virus-free medium was added back to the cells. For selection, 1.3 mg/ml G418 was used. After selection procedure, expression of transgene was validated by western blot.

### Western blots

For western blot analysis, cells were harvested using trypsin and washed 2x with PBS. Cells were resuspended in an appropriate volume of lysis buffer (50 mM Tris-HCl pH 7.4, 1 mM EGTA, 1 mM EDTA, 1% Triton X100, 0.27 M sucrose, 1mM DTT, cOmplete EDTA-free protease inhibitor (Roche)) and incubated for 30 min on ice. Lysates were cleared by centrifugation and protein concentration was determined by Bradford (Biorad). 10 µg of protein was boiled for 5 min at 95°C and then loaded on a 4-15% SDS gel (Biorad).

Proteins were transferred on a nitrocellulose membrane by wet blotting (Biorad) at 120V for 60 min. Membranes were blocked with 3% non-fat dry milk in TBS + 0.05% Tween-20 (blocking buffer) for 1h at RT. Primary antibodies were added for overnight incubation at an appropriate dilution (see below). The next day membranes were washed 3x with TBS + 0.05% Tween-20 (TBST) and incubated with the secondary antibody diluted in blocking buffer. After 3 washes with TBST, membranes were developed with ECL Clarity (for GAPDH) or Super Signal West femto chemiluminescence substrate (Thermo Fisher). Images were taken with ChemiDoc Touch system (Bio-Rad).

Antibodies dilutions used for western blot: rabbit anti AZI2/NAP1 (Abcam 1:1000), mouse anti NBR1 (Abnova 1:1000), mouse anti GAPDH (Sigma, 1:25,000), mouse anti p62 (BD Bioscience, 1:1000), rabbit anti p62 pSer349 (Cell Signaling, 1:1000), rabbit anti p62 pSer403 (Cell Signaling, 1:1000), rabbit anti TAX1BP1 (Cell Signaling, 1:1000). Secondary antibodies: goat anti rabbit HRP (Jackson Immunoresearch, 1:10,000), goat anti mouse HRP (Jackson Immunoresearch, 1:10,000).

For quantification, ImageJ was used. A rectangle was drawn around the corresponding protein band and the intensity was plotted using the “Plot Lanes” function. Measured values were adjusted and corrected by GAPDH as a loading control.

### Live cell imaging

Cells were seeded in 10w glass bottom imaging chambers (Greiner Bio one) at 6000 cells/ well. 48h later cells were treated with drugs and either imaged using a Visitron Live Spinning Disk microscope (Plan-Apochromat 63x/1.4 Oil DIC objective and an EM-CCD camera; including a temperature & CO2 controlled environment). Standard imaging settings were a gain of 500 for all channels and 100 ms exposure for GFP & mScarlet and 200 ms for iRFP. Laser settings were the following: 5% laser power for GFP, 10% laser power for mScarlet and 10% laser power for iRFP. For dox-inducible TAX1BP1 constructs, iRFP settings were a gain of 500, 150 ms exposure and 15% laser power. Settings were kept consistent across replicates. Multiple areas within the wells were imaged to obtain comparable numbers of cells.

For quantifications a macro for ImageJ was used as described before (Turco *et al*., 2021). In short, binary pictures were generated by thresholding. Analyze punctae with a cut off of “>3 pixel units” was used to count puncta and to save coordinates as ROIs. ROIs from one channel were overlayed with the other channel(s) to determine the degree of colocalization. Threshold was determined manually and was kept consistent across replicates. Cells with a significant amount of background noise were excluded from the analysis

### Immunofluorescence & confocal microscopy

Immunofluorescence was performed similar to previous studies (Turco *et al*., 2021). In short, cells were seeded in 6 well plates on cover slips. After 24h the cells were treated, washed 3x with PBS and fixed with 4% Paraformaldehyde in PBS for 20 min at RT. After fixation, the cells were washed 3x with PBS and permeabilized with 0.1% Triton X100 in PBS for 5 min at RT except for stainings that included FIP200. For FIP200, cells were permeabilized with 0.25% Triton X100 in PBS for 15 min at RT. Cells were washed 2x with PBS, incubated with blocking buffer (1% BSA in PBS) at RT for 1h and incubated with the primary antibodies for 1h at RT in a humid chamber except for FIP200. For FIP200, coverslips were incubated immediately with the primary antibody in a humid chamber for 1h at 37°C without a previous blocking step. The cells were washed 3x with PBS and incubated with the secondary antibody for 1h at RT in a humid chamber. Cells stained with FIP200 were incubated for 1h at 37°C in a humid chamber with the secondary antibody. All cells were washed 3x with PBS and mounted on glass slides (Roth) by transferring them onto a droplet of the mounting medium DAPI-Fluoromont-GTM (Southern Biotech).

Antibody dilutions (in blocking buffer): rabbit anti ATG9 (1:100, Cell Signaling), rabbit anti FIP200 (1:200, Cell Signaling), mouse anti p62 (1:100, BD Bioscience), rabbit anti AZI2/NAP1 (1:100, Abcam), rabbit anti TAX1BP1 (1:100, Cell Signaling), anti pTBK1-S172 (1:100, Cell Signaling), goat anti rabbit Alexa Fluor 546 (1:1000, Invitrogen), goat anti mouse Alexa Fluor 488 (1:1000, Invitrogen). Images were taken with a Zeiss LSM 700 confocal microscope equipped with Plan-Apochromat 63x/1.4 Oil DIC objective. To avoid cross-contamination between fluorochromes, each channel was imaged sequentially using the multitrack recording module before merging. Images from fluorescence and confocal acquisitions were processed and analyzed with ImageJ software.

For quantifications a macro for ImageJ was used as described before (Turco *et al*., 2021). In short, binary pictures were generated by thresholding. Analyze puncta with a cut off of “>3 pixel units” was used to count the number of puncta and to save coordinates as ROIs. ROIs from one channel were overlayed with the other channel(s) to determine the degree of colocalization. Threshold was determined manually and was kept consistent across replicates. Cells with a significant amount of background noise were excluded from the analysis

### Protein production

N-domain (aa420-506 of TAX1BP1), GFP-N-domain, Tetra-ubiquitin (4xUB), GFP and GFP-CC2-LIR1 were cloned into a pGEX -4T1 plasmid vector backbone. mCherry-p62 S403E was cloned into the pET vector backbone. After transformation into competent Rosetta *E. coli*, a single colony was picked and incubated overnight in 100 ml of LB including the appropriate selection marker. The next day, the cultures were scaled up to the range of liters. The cultures were incubated at 37°C until an OD_600_ of 0.6 was reached and then the expression was induced by addition of 0.2 mM isopropylthiogalactoside (IPTG). The cells were harvested after 18-20h at 18°C and pellets were flash-frozen in liquid nitrogen and stored at −70°C. For mCherry-p62 S403E, the cells were incubated until they reached the OD 0.8. The expression was induced with 0.2 mM IPTG and the cells were harvested after 5h shaking at 25°C, flash-frozen and stored at −70°C.

10x-His-GFP-TAX1BP1, 10x-His-GFP-TAX1BP1ΔN, 10x-His-GFP-TAX1BP1ΔZnF, 10x-His-mCherry-TAX1BP1, 10x-His-mCherry-TAX1BP1ΔN, GST-TAX1BP1, GST-TAX1BP1ΔZnF, GST-TAX1BP1ΔN, GST-TAX1BP1 L442A, GST-TAX1BP1 Y449A were cloned into the pLIB vector backbone for insect cell culture expression. Codon optimized GFP-TBK1, TBK1, SINTBAD and GFP-SINTBAD were generated by gene synthesis (GenScript) for insect cell expression. The plasmids were transformed into electrocompetent DH10BacY cells. For the transformation, 1 µl of the plasmids at a concentration of 25 ng/µl were mixed with bacteria and electroporated using a Biorad Micropulser at the appropriate settings for bacteria (2.5 kV potential difference). For recovery, the electroporated cells were mixed with 5 ml of LB and incubated for at least 5h at 37°C. Afterwards, the cells were plated on LB agar plates containing 50 µg/ml Kanamycin, 7 µg/ml Gentamicin, 10 µg/ml Tetracycline, 200 µg/ml X-Gal and 165 µM IPTG. White colonies were screened on the next day by colony PCRs. Positive colonies were picked and incubated with 10 ml of LB & appropriate antibiotics over night at 37°C. The next day the bacmids were isolated and the DNA measured. The bacmids were used freshly for insect cell transfections. 1 million of Sf9 insect cells were seeded on 6 well plates. 5 µg of DNA and 5 µl of FuGene HD was used for transfection. The expression of the reporter YFP was tracked over several days by microscopy. Once most cells showed YFP expression (usually after 5 days), the cell suspension (V0) was added to 30 ml of insect cells at 1 mio/ml. Viability of cells and YFP expression was monitored for several days. Once, the cells showed viability drop to below 90%, the cells were spun down and the supernatant was filtered with a 0.45 µm filter and stored at 4°C. For expression 1 ml of the V1 was added to 900 ml of Sf9 insect cells at 1 mio/ml. Once the viability dropped below 90%, the cells were harvested, washed once with PBS, flash frozen and stored at −70°C GFP-NBR1 and NBR1 were cloned into the pCAGGSsH-C vector and expressed in HEK293 via the VBCF ProTech facility. Pellets were stored at −70°C.

### Protein purification

mCherry-p62 S403E, GST-4xUB, GFP-NBR1, NBR1, GFP-TAX1BP1 FL, mCherry-TAX1BP1 FL were purified as described previously (Turco *et al*., 2021; Zaffagnini *et al*., 2018).

Truncations and point mutants of TAX1BP1 were purified following the same procedure as for the full-length TAX1BP1 (FL) protein.

GFP-TBK1, TBK1, SINTBAD & GFP-SINTBAD were also purified as previously described (Adriaenssens *et al*., 2023).

GST-N domain, GFP-N domain, GST-GFP-CC2-LIR1 were purified from bacterial pellets. Pellets corresponding to 2 L culture were resuspended in lysis buffer (50 mM HEPES, pH 7.5, 300 mM NaCl, 1 mM DTT, 2 mM MgCl_2_, cOmplete EDTA-free protease inhibitor (Roche), DNase (Sigma),) and lysed by sonication. To clear the lysate, the suspension was centrifugated with 18 000 rpm at 4°C. The supernatant was filtered through a 0.45 µm filter and incubated with 4 ml pre-equilibrated GST beads (GE Healthcare). After 1h incubation rolling at 4°C, the beads were washed 5x with Wash Buffer I ((50 mM HEPES, pH 7.5, 300 mM NaCl, 1 mM DTT), once with Wash Buffer II (50 mM HEPES, pH 7.5, 700 mM NaCl, 1 mM DTT) and again 2x with Wash Buffer I. For elution, beads were either resuspended in 3 ml Wash Buffer I + TEV protease for cleavage overnight or by incubation with elution buffer (Wash Buffer I + 50 mM glutathione (Sigma) -pH adjusted to 7.5). The next day, the eluate was filtered through a 0.45 µm filter, concentrated to 500 µl using an appropriate protein concentrator (Millipore) and further purified by size exclusion chromatography on a Superdex 200 Increase 10/300 column (GE Healthcare) in 25 mM HEPES—pH 7.5, 150 mM NaCl, 1 mM DTT.

### Microscopy-based protein interaction assay

Microscopy-based interaction assay was performed as previously described (Sawa-Makarska *et al*, 2020; Turco *et al*., 2021; Zaffagnini *et al*., 2018). In short, GST (GE Healtcare), GFP-trap or RFP-trap beads (both Chromotek) were washed first 2x with H2O and then equilibrated 2x with SEC buffer (25 mM HEPES, pH 7.5, 150 mM NaCl, 1 mM DTT). Beads were resuspended in 40 µl of SEC buffer and the bait protein was added. For experiments with GST-TAX1BP1 (FL, domain, truncations, point mutants) 2.5 µM of bait protein was added to the beads. The beads were incubated for 1h at 4°C and washed afterwards 3x with SEC Buffer. The beads were pipetted into a 384 well plate containing 1 µM GFP-NBR1 as prey in a total volume of 20 µl. For experiments with GST-4xUB, 5 µM bait protein was added and treated as described above. Beads were pipetted into 1 µM GFP-TAX1BP1 (FL, truncation) prey solution. For experiments with the GABARAP competition, GFP-trap beads were incubated with 2.5 µM GFP-NBR1 and treated as described above. Beads were added to 500 nM mCherry-TAX1BP1 and increasing amounts of GABARAP protein (0 nM – 500 nM – 1 µM – 5 µM). For experiments in Fig. 7G, GSH beads were first incubated with a mix of 4.5 µM GST + 0.5 µM GST-4xUB as a bait for 1h. After the incubation, the beads were washed to remove the unbound proteins and further incubated with 1 µM mCh-p62 and 500 nM NBR. After 30 min of incubation, 500 nM GFP-TAX1BP1 FL or GFP-TAX1BP1ΔZnF, together with 5 µM GABARAP, was added to the beads. Prey & bait were incubated for 15 min to 30 min at RT and imaged imaged with a Zeiss LSM 700 confocal microscope equipped with Plan Apochromat 20X/0.8 WD 0.55 mm objective. For each experimental condition three biological replicates were performed. For the quantification, we employed an artificial intelligence (AI) script that automatically quantifies signal intensities from microscopy images by drawing line profiles across beads and recording the difference between the minimum and maximum grey values along the lines. The AI was trained to recognize beads employing cellpose (Stringer *et al*, 2021). Source code available at https://www.maxperutzlabs.ac.at/research/facilities/biooptics-light-microscopy. Background corrected values were plotted and subjected for statistical analysis. For GABARAP competition, we normalized to the control (0 nM GABARAP) to minimalize the inter-experiment variability.

### Pulldowns

For pulldowns with cell lysates, HeLa PentaKO cells were seeded at 500k/well. 24h later the cells were transfected with plasmids encoding GFP-NBR1 (FL, truncations, point mutants). 48h after transfections, cells were harvested and lysed using lysis buffer (50 mM Tris-HCl pH 7.4, 1 mM EGTA, 1 mM EDTA, 1% Triton X100, 0.27 M sucrose, 1mM DTT, cOmplete EDTA-free protease inhibitor (Roche)). GST-TAX1BP1 coated beads were prepared as described above. 100-200 µg of total protein lysates was mixed with beads and incubated for 1h at 4°C. After the incubation, beads were washed 3x with IP wash buffer (20 mM Tris-HCl pH 7.4, 150 mM NaCl, 1 mM EDTA, 0.05% Triton-X100, 5% glycerol) to remove unbound material. 40 ul of 2x loading dye was added to the beads. Samples were boiled for 10 min at 95°C. SDS-Page and western blotting for detection was performed as described above.

### Condensate assay

*In vitro* condensation assay has been performed as previously described (Turco *et al*., 2021; Turco *et al*., 2019; Zaffagnini *et al*., 2018). In short, proteins were spun down with max speed for 10 min at 4°C. 2 µM mCherry-p62 S403E, 1 µM NBR1, 2 µM GFP-TAX1BP1 (FL, ΔN or ΔZnF) were mixed in SEC buffer (25 mM HEPES, pH 7.5, 150 mM NaCl, 1 mM DTT) in a 384 well plate. Buffer without NaCl was used to keep the salt level at 150 mM NaCl. 5 µM GST-4xUB was added at the microscope to trigger condensation of the proteins. Images were taken as a time course over 2h with a Spinning disk microscope equipped with LD Achroplan 20X/0.4 Corr objective and EM-CCD camera.

For the reconstitution of the recruitment of TBK1 & SINTBAD to p62 condensates, 500 nM mCherry-p62 S403E, 250 µM NBR1, 250 nM mCherry-TAX1BP1, 250 nM SINTBAD, 500 nM GFP-TBK1, 500 nM GFP-SINTBAD were mixed in SEC buffer (25 mM HEPES, pH 7.5, 150 mM NaCl, 1 mM DTT) in a 384 well plate. 1.5 µM GST-4xUB was added to at the microscope to trigger condensation of the proteins.

For the competition assay with the GABARAP proteins, 500 nM mCherry-p62 S403E, 250 nM NBR1, 250 nM GFP-TAX1BP1 and increasing amounts of GABARAP (0 nM – 500 nM – 1 µM – 5 µM) were mixed in SEC buffer (25 mM HEPES, pH 7.5, 150 mM NaCl, 1 mM DTT) in a 384 well plate. 1.5 µM GST-4xUB was added to at the microscope to trigger condensation of the proteins.

For quantifications, the same macro was used as previously described (Turco *et al*., 2021; Zaffagnini *et al*., 2018). In short, rolling ball background subtraction of ImageJ was performed, followed by thresholding and puncta counting across all channels and timepoints using the Analyze puncta function.

### Peptide Array

The peptide array was synthetized by the Peptide Synthesis facility of the IMP (https://cores.imp.ac.at/peptide-synthesis/). The entire NBR1 protein sequence was spotted as 12 aa/spot in a 2 aa window. The membrane was first soaked in methanol and then washed 3x with TBS. Membrane was incubated with blocking buffer (3% BSA dissolved in 1xTBS) overnight at 4°C.The next day the membrane was washed 1x with TBST (TBS + 0.1% Tween20) and incubated with our protein (GFP-TAX1BP1) at a concentration of 50 ng/ml in 15 ml of SEC Buffer (25 mM HEPES, pH 7.5, 150 mM NaCl, 1 mM DTT) for 2h at RT. After the incubation, the membrane was washed 3x with TBST.

For detection a western blot using a PVDF membrane was performed. After the transfer by wet blotting, the membranes were treated as described above.

### Statistical analysis

For the statistical analysis Prism GraphPad was used. For all microscopy-based interaction assays a one-way ANOVA was performed. For immunofluorescence images student’s t-test was performed. For live cell imaging data, two-way ANOVA or student’s t-test was performed. For western blot quantifications one-way ANOVAs was performed. For condensation assays, either one-way ANOVA or student’s t-test was performed depending on the number of conditions. For comparing the number of foci between TAX1BP1 WT and ΔZnF we performed a one sample t and Wilcoxon test. p values were corrected for multiple testing using the suggested method. Statistical significance is indicated with *P<0.05, **P<0.005, ***P<0.001, ****P<0.0001, ns = not significant. Error bars are reported as mean ± standard deviation.

## Acknowledgments

We thank all members of the Martens lab for fruitful discussions and Susanna Tulli and Elias Adriaenssens for comments on the manuscript. We also thank Elias Adriaenssens for providing SINTBAD protein, Michael Lazarou for providing the PentaKO HeLa cells, Jean-Marc Furlano for help with the NBR1 – TAX1BP1 interaction assays, Lea Radzuweit for help with the purification of mCherry-p62 S403E and the Clausen lab (IMP, Vienna) for help with the peptide array. We further thank the Max Perutz Labs BioOptics facility and the Mass Spectrometry facility for technical support. Proteomics analyses were performed using the VBCF instrument pool. We also thank the VBCF Protech facility for the HEK cell expressions.

## Funding

Austrian Science Fund (FWF P30401-B21, PAT7165623, W1261 and F79)

European Union’s Framework Programme for Research and Innovation Horizon 2020 (2014-2020) under the Marie Curie Skłodowska Grant Agreement Nr. 847548

## Competing interests

Sascha Martens is member of the scientific advisory board of Casma Therapeutics. The other authors declare no competing interests.

## Supplementary Figures

**Fig. S1:**
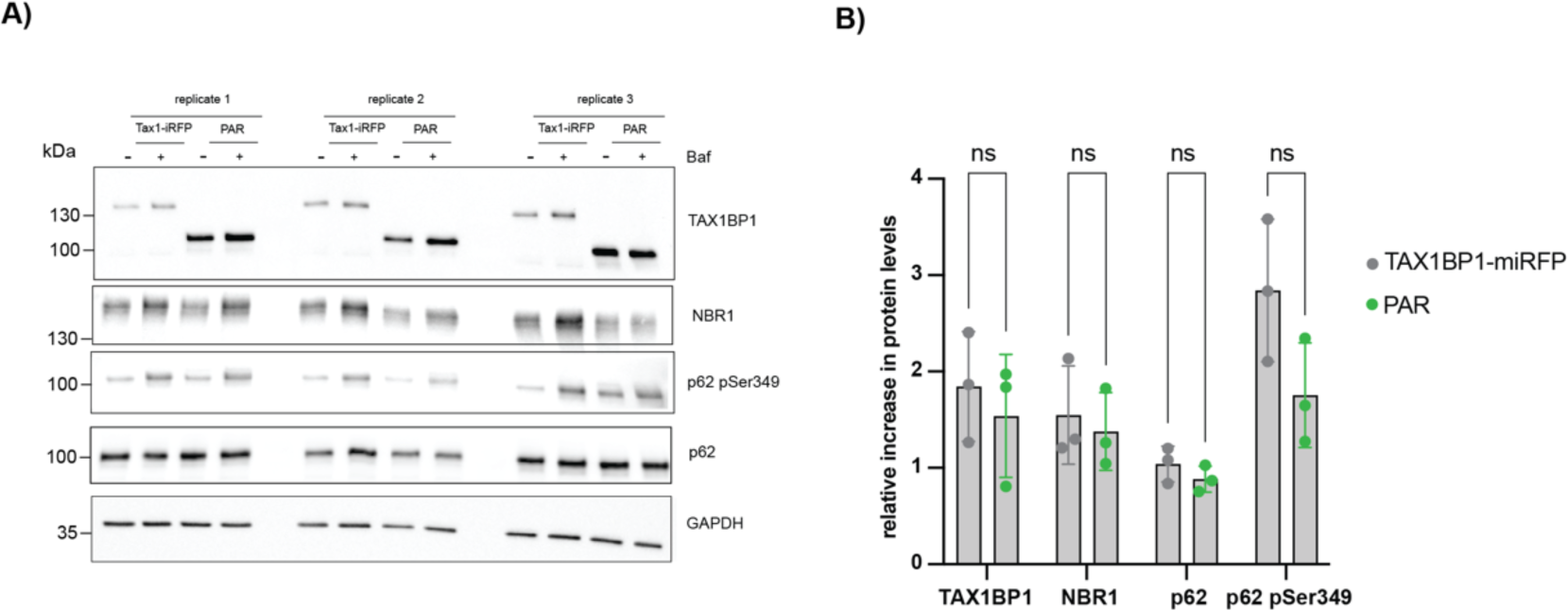
TAX1BP1 is not a constitutive component of p62 condensates in cells. A) Western blot for the validation of the CRISPR knock-in cell line. B) Quantification of A. The increase between untreated and Bafilomycin treatment is plotted and compared to the parental cell line (PAR). Data in B shows the mean +/− s.d. from three independent experiments. Two-way ANOVA with Sidak’s multiple comparison test was performed in B. ns = not significant

**Fig. S2:**
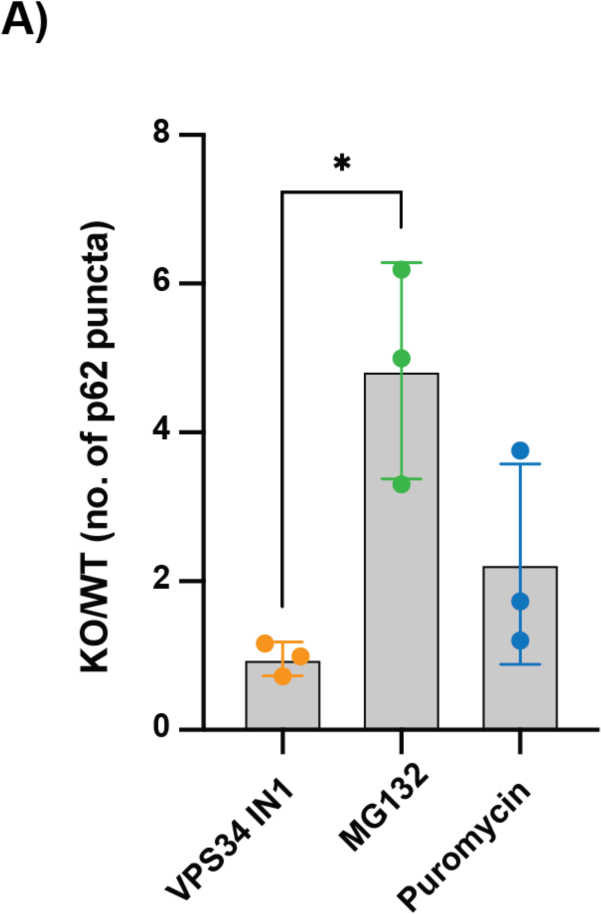
TAX1BP1 KO cells accumulate more p62 condensates upon proteasomal inhibition. A) Quantification of the relative increase in the number of p62 puncta between the indicated treatments. Data in A shows the mean +/− s.d. from three independent experiments. One-way ANOVA with Tukey multiple comparison test was performed in B. *P<0.05, ns = not significant

**Fig. S3:**
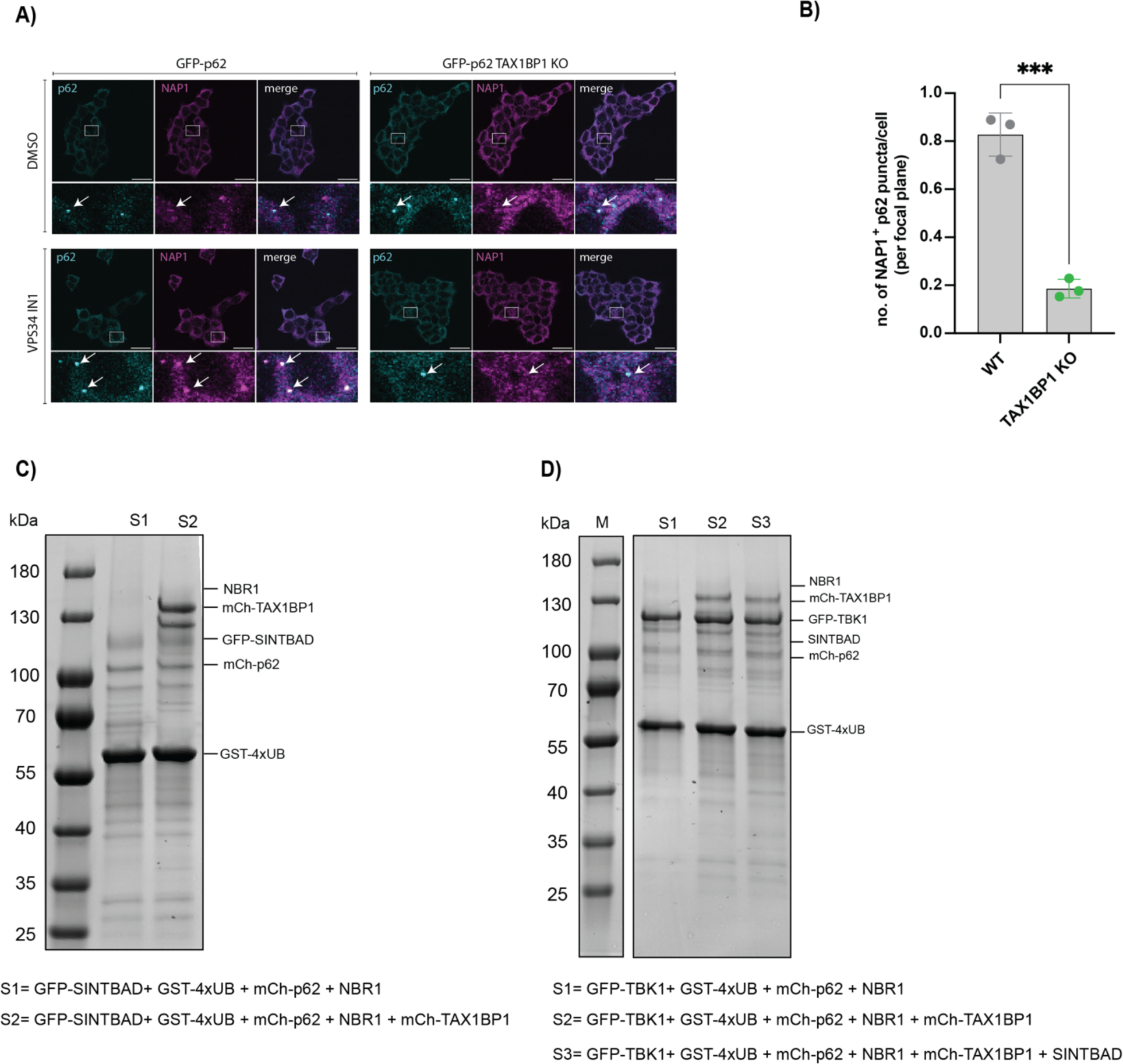
TAX1BP1 recruits TBK1 to p62 condensates via the adapter protein SINTBAD. A) Representative immunofluorescence images of cells stained for p62 and NAP1 untreated and upon autophagy inhibition (scale bar = 20 µm). B) Quantification of the number of p62 puncta positive for NAP1 upon autophagy inhibition. C) SDS Page gel as loading control for condensation assay in 3G D) SDS Page gel as loading control for condensation assay in 3H (Protein bands are labelled). Data in B shows the mean +/− s.d. from three independent experiments. Unpaired t-test was performed in B. ***P<0.001.

**Fig. S4:**
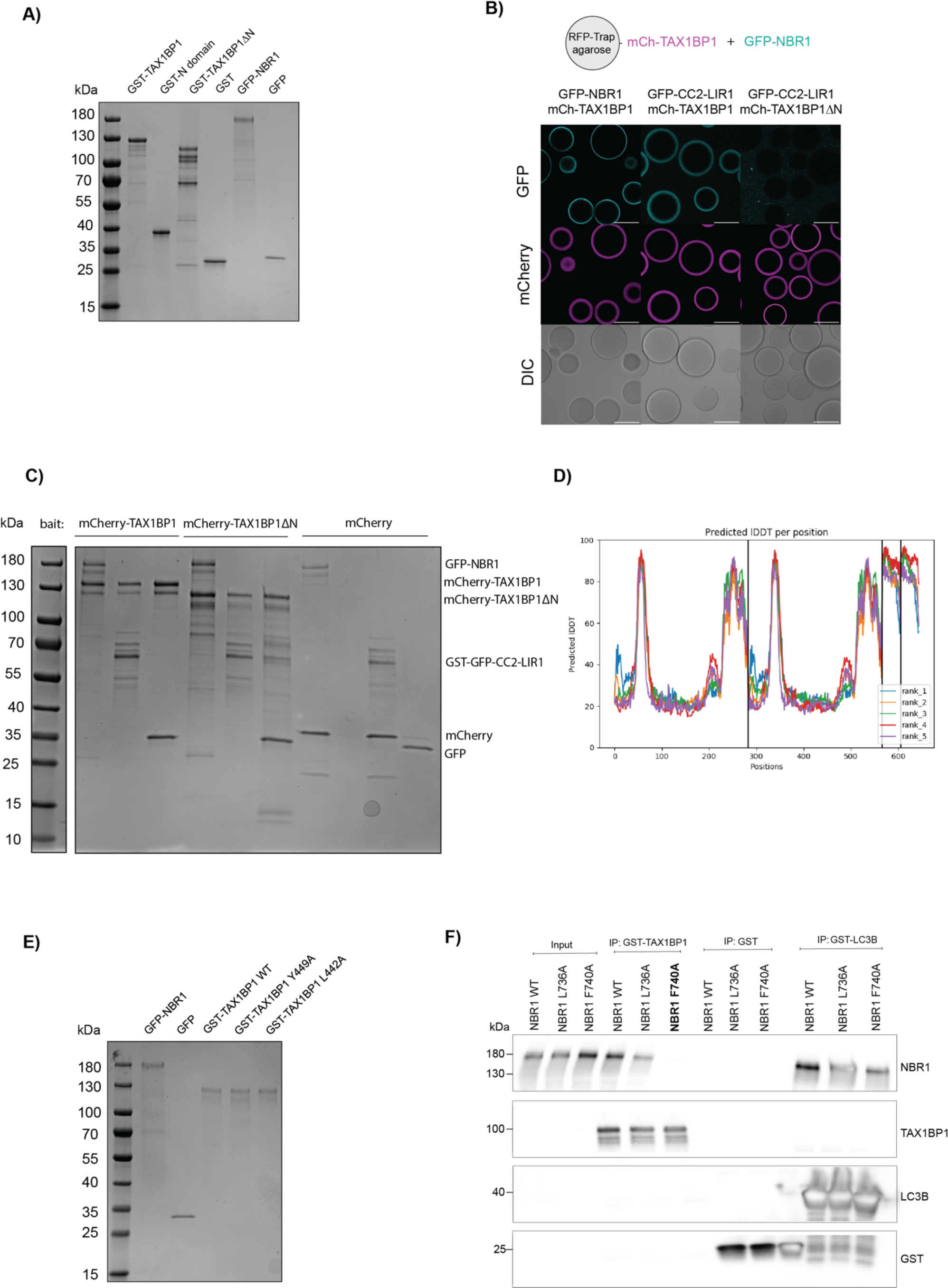
Mapping the interaction between TAX1BP1 and NBR1. A) SDS Page gel as loading control for microscopy-based interaction assay in 4A. B) Representative images for microscopy-based interaction assay with GFP tagged NBR1 fragment (CC2-LIR2) and mCherry-TAX1BP1 FL or ΔN on RFP-trap beads (scale bar = 80 µm). C) SDS Page gel as loading control for interaction assay from S4B. D) pLDDT plot for the AlphaFold 2 prediction in 4E. E) SDS Page gel as loading control for interaction assay in 4F. F) Western blot of pulldown with cells expressing NBR1 point mutants including GST-LC3B as bait to determine the effect of point mutant on GABARAP and LC3B binding.

**Fig. S5:**
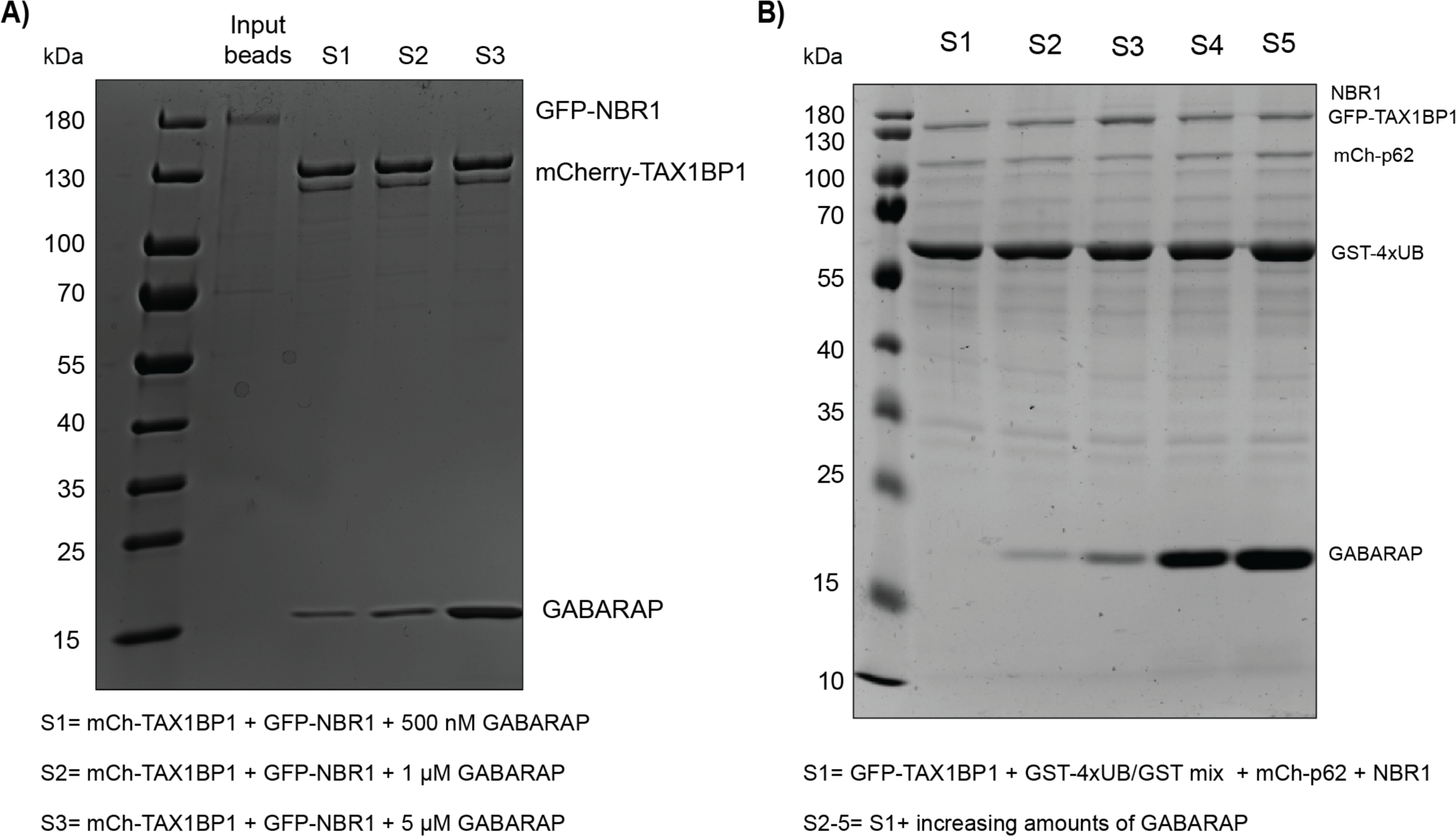
GABARAP competes with TAX1BP1 for NBR1 binding. A) SDS Page gel as loading control for interaction assay in 5A. B) SDS Page gel as loading control for GABARAP competition in condensation assay in 5E.

**Fig. S6:**
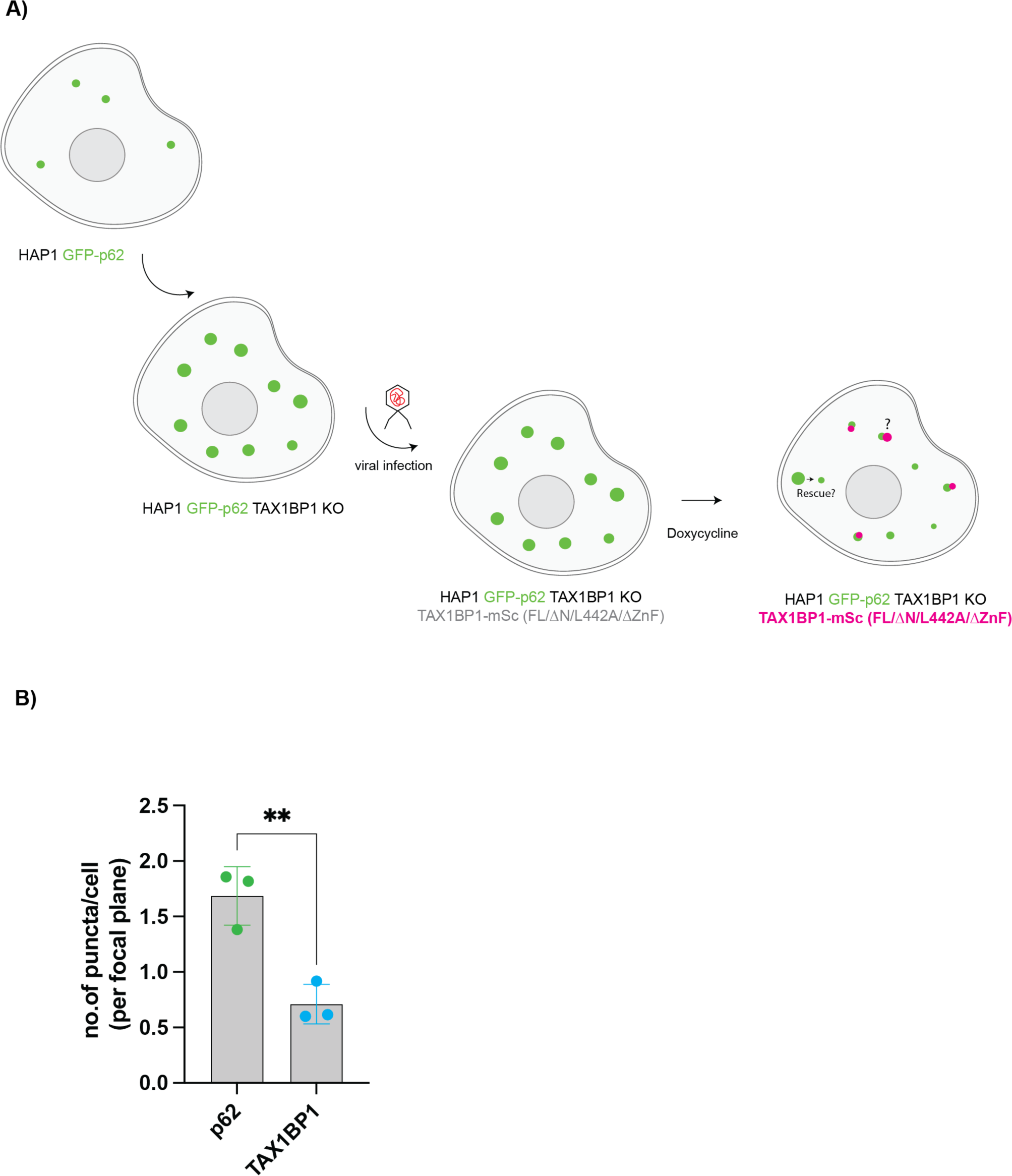
The TAX1BP1 – NBR1 interaction is required for the autophagic flux of p62 condensates. A) Schematic overview for the experimental set-up of the experiment shown in Fig. 6. B) Quantification of the number of p62 and TAX1BP1 foci in R-WT cells upon VPS34 IN1 treatment. Data in B shows the mean +/− s.d. from three independent experiments. Unpaired t-test was performed in B. **P<0.005.

**Fig. S7:**
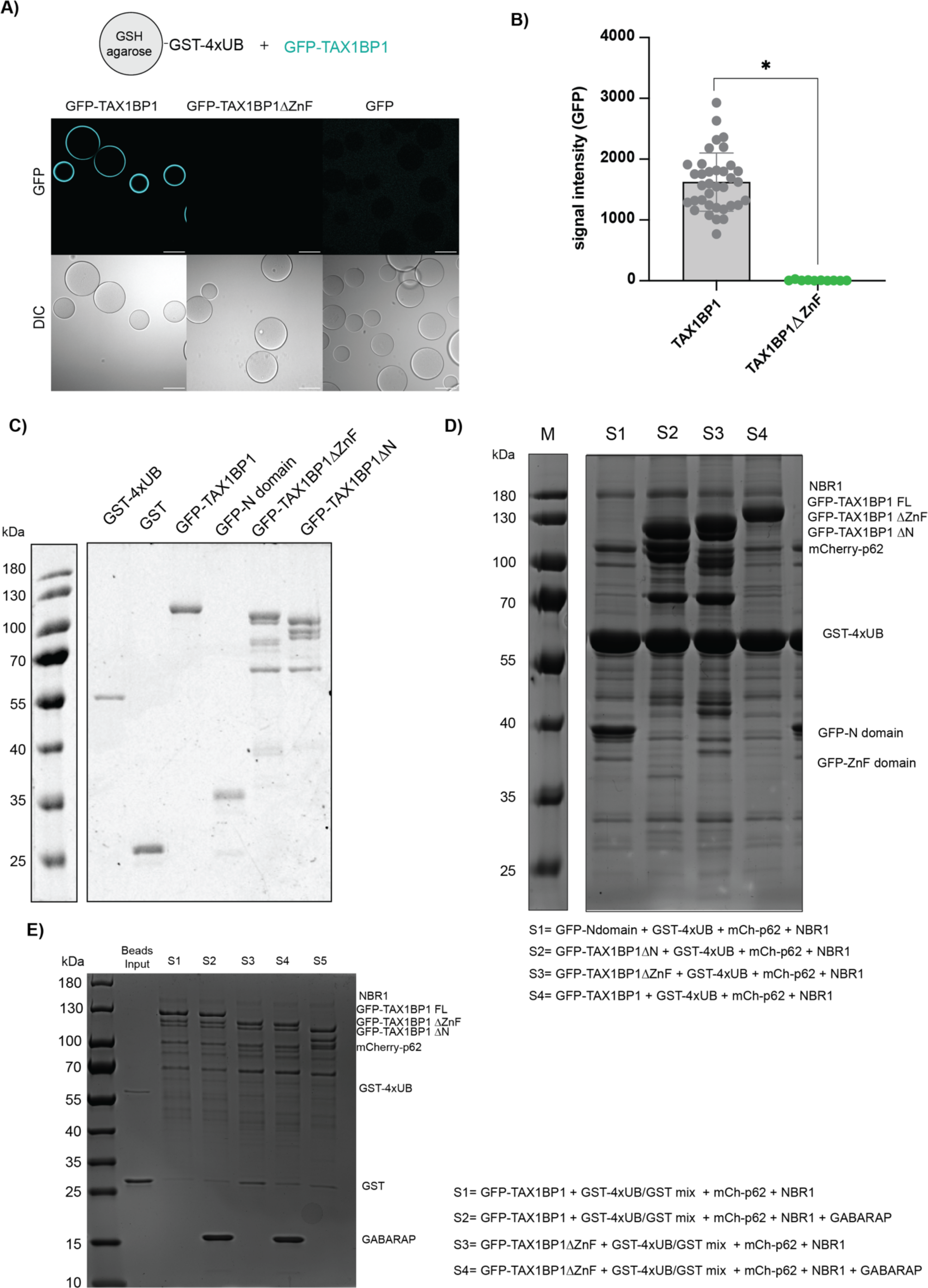
Ubiquitin interaction stabilizes the recruitment of TAX1BP1. A) Representative images of a microscopy-based protein-protein interaction assay using the indicated proteins (scale bar = 80 µm). B) Quantification of S7A. C) SDS Page gel as a loading control for the microscopy-based interaction assay shown in S7A. D) SDS Page gel as a loading control for the condensation assay shown in 7A. E) SDS Page gel as a loading control for the microscopy-based interaction assay in 7G. Data in B shows the mean +/− s.d. from three independent experiments. Each data point in B represents the mean signal intensity for an individual bead. Nested one-way ANOVA with Dunnett’s multiple comparison test was performed in B. *P<0.05.

